# Encoding neural representations of time-continuous stimulus-response transformations in the human brain with advanced deep neural networks

**DOI:** 10.1101/2025.09.19.677324

**Authors:** Sabine Haberland, Hannes Ruge, Holger Frimmel

## Abstract

Human behavior arises from the continuous transformation of sensory input into goal-directed actions. While existing analytical methods often break time into discrete events, the stages and underlying representations involved in stimulus-response (S-R) transformations within time-continuous, complex environments remain only partially understood. Encoding models, combined with deep neural networks (DNNs) for feature generation, offer a promising framework for capturing these neural processes. While DNNs continue to improve in performance, it remains unclear whether these advances translate into more accurate models of brain activity. To address this, we collected fMRI data from participants (N = 23) as they played arcade-style video games and applied DNN-based encoding models to predict voxel-level brain activity. We compared the prediction accuracy of features from three DNNs at different stages of development within our encoding model. We show that the most advanced DNN provides the most predictive feature space for neural responses, while also exhibiting a closer hierarchical alignment between its internal representations and the brain’s functional organization. These results enable a more fine-grained characterization of time-continuous S-R transformations in high-dimensional visuomotor tasks, progressing along the dorsal visual stream and extending into motor-related regions. This approach highlights the potential of machine learning to advance cognitive neuroscience by enhancing the ecological validity of experimental tasks.

## 1 Introduction

The human brain is capable of rapidly processing complex, dynamic information of the environment and responding flexibly with appropriate actions, whether it is catching a flying ball or navigating through traffic. For decades, the study of these underlying neural processes has been a central focus of research and a key source of inspiration for the development of artificial neural networks in machine learning. Even though neuroscience and machine learning have advanced rapidly in recent decades, it is only recently that the two fields have begun to exchange concepts and benefit from each other’s progress (Naselaris et al., 2018). These brain-inspired neural networks are now being used as computational tools to better understand neural processes themselves (Cichy & Kaiser, 2019; Kriegeskorte, 2015). While machine learning has primarily focused on improving performance to automate complex tasks, it remains uncertain whether these advancements can be translated back into neuroscience (Botvinick et al., 2020).

Factorial experimental designs, which are constrained by a low-dimensional stimulus space and trial-based structure, have provided valuable insights into the neurofunctional architecture of the human brain. However, they may not fully capture the complexity of real-world processing, which limits their ecological validity. The encoding model approach has emerged as a promising method to overcome these limitations (van Gerven, 2017). Using this method, prior studies have modeled neural activity patterns evoked by complex stimuli, such as naturalistic images, using high-dimensional stimulus feature spaces including hand-crafted features like a Gabor wavelet pyramid (GWP) basis (Kay et al., 2008).

A breakthrough has been achieved by combining encoding models with deep neural networks (DNNs). In this approach, the activations of the DNNs serve as nonlinear representations of stimulus features for the encoding model. Studies have shown that convolutional neural networks (CNNs), which achieve human-level accuracy in object classification, can be used to generate suitable stimulus features. This enables a more precise analysis of voxel activity along the ventral visual stream when subjects are presented with naturalistic images than a GWP model (Güclü & van Gerven, 2015). Additionally, studies have revealed a hierarchical structure in these representations, suggesting that the layered architecture of the DNNs captures the stages of visual processing and complexity observed in the human brain along the ventral visual stream (Cichy et al., 2016; Eickenberg et al., 2017; Güclü & van Gerven, 2015). In these experiments, subjects passively viewed visual stimuli and were not required to perform specific actions. This approach allows the investigation of the ventral visual stream, which extends from the primary visual cortex in the occipital lobe to the temporal lobe. The ventral stream is primarily responsible for generating detailed visual representations, which are important for object recognition and categorization (Goodale, 2014; Grill-Spector & Weiner, 2014). In contrast, the dorsal visual stream serves as a second pathway, extending from the primary visual cortex to the posterior parietal lobe (Goodale & Milner, 1992). Studies have demonstrated that this stream is activated when visual information must be transformed into motor actions (Goodale, 2011). Moreover, recent studies have shown that stimulus-response (S-R) transformations in visuomotor tasks involve functional connectivity between the posterior parietal cortex (PPC) and the premotor cortex (PMC) (Brovelli et al., 2015; Creem-Regehr, 2009; Kravitz et al., 2011). Along the dorsal stream, studies have also demonstrated a hierarchical similarity between stimulus processing in the brain and representations in DNNs during action recognition (Güclü & van Gerven, 2017a). Due to technical limitations, the dorsal visual stream has remained underexplored, and its neural representations are less understood than those in the ventral stream. This is partly because sensorimotor integration involves complex, dynamic functions that are hard to capture with traditional neuroimaging and often require elaborate behavioral paradigms. Recent advances in machine learning, however, provide new ways to study these processes.

These advancements, particularly the combination of deep learning with reinforcement learning (RL), have led to the development of deep Q-networks (DQNs). DQNs can solve tasks involving complex, time-continuous stimuli that require motor responses at or above human-levels (Mnih et al., 2015). A DQN approximates a Q-function, which estimates the expected cumulative discounted reward for taking an action in a given state, aiming to maximize the total reward over time. When DQNs are integrated into a DNN-based encoding model, this approach has successfully predicted human behavior during arcade gameplay (Haberland et al., 2025; Mohr et al., 2019). It also provides ways to explore functional similarities and model neural processes underlying S-R transformations in visuomotor tasks, as demonstrated by studies that recorded brain activity in participants playing video games (Cross et al., 2021). These results suggest the human brain may use similar computational principles to interact in complex, dynamic environments.

In recent years, the development of DQNs has progressed significantly and enabled them to achieve superhuman performance across various tasks (Espeholt et al., 2019). At the behavioral level, it has already been demonstrated that advancements in machine learning can be leveraged to improve prediction accuracy when modeling human behavior in complex, time-continuous experimental tasks at a fine-grained temporal scale (Haberland et al., 2025). Building on previous findings, we aimed to investigate whether advancements in machine learning can capture neural representations underlying the steps of S-R transformations in complex, time-continuous tasks, thereby enabling a more detailed characterization and localization of these processes.

Therefore, we analyzed fMRI data from *N* = 23 participants, each measured on three separate days while playing the arcade games Breakout, Space Invaders, and Enduro. The three games represent different gameplay mechanics. We applied a DQN-based encoding model approach to predict the underlying fMRI activation patterns. To assess progress in RL, we evaluated two recently developed DQNs, ApeX (Horgan et al., 2018) and SEED (Espeholt et al., 2019), and a baseline DQN (Mnih et al., 2015), as feature-generating mappings within an encoding model, comparing their prediction accuracy for voxel activity. While all three DQNs are capable of learning to play arcade games after sufficient training, they differ in their architectures and training methods. The baseline DQN is a standard feed-forward CNN. Although Ape-X and SEED share structural similarities with the baseline DQN, they incorporate additional architectural components and significantly enhance data generation through parallel sampling and optimized hardware utilization. Ape-X and SEED also integrate a dueling architecture and SEED further introduces a long short-term memory (LSTM), a recurrent neural network (RNN) that incorporates past experiences into decision-making (Hochreiter & Schmidhuber, 1997). These advancements result in a significant increase in performance that surpasses human-level performance in all three games.

Using the DQN-based encoding model, we replicated previous findings showing that this method is well-suited for modeling S-R transformations in time-continuous and complex environments. Building on these results, we confirm our hypothesis that recent advances in machine learning enable a more fine-grained characterization of neural representations, particularly in higher-level cognitive control regions, making this approach even more valuable for future investigations of temporally structured behavior. By comparing the spatial representations in the brain with those in the DQNs, we uncover a hierarchical correspondence between DQN layers and visuomotor processing stages in the dorsal stream during visuomotor tasks. These findings highlight the potential of machine learning to advance neuroscience.

## 2 Methods

### 2.1 Participants

The participants (*N* = 23, age range: 19 − 35, 9 female, 14 male) were right-handed, neurologically healthy, and had normal or corrected-to-normal vision. They were recruited from the population of the TUD Dresden University of Technology. Due to incomplete data collection from one participant, only *N* = 22 datasets were available for the Space Invaders task. None of the participants had participated in our first behavioral experiment (Haberland et al., 2025). They were compensated with 10 EUR per hour, along with an additional variable bonus based on their game score (range: 3.87 EUR - 15.98 EUR per game).

### 2.2 Experimental design and task

#### 2.2.1 Ethics statement

The experimental protocol was approved by the Ethics Committee of the TUD Dresden University of Technology (EK 359072019) and was conducted in accordance with the Declaration of Helsinki. All participants were informed about the study’s objectives and procedures and provided written informed consent before participation.

#### 2.2.2 Experimental paradigm and procedure

Each participant attended three separate appointments on three different days. On each day, one of the three arcade games, Breakout, Space Invaders, or Enduro, was played. The games were selected in a pseudo-random order. Each appointment began with the participants reading the gaming instructions for the chosen game, including finger placement, controls, objectives, and scoring. Then they trained the game for 35 minutes in a mock MRI scanner to become familiar with the environment and the game. After the training, participants transitioned to the MRI scanner for the main experiment. Following a T1-weighted high-resolution anatomical scan, a field map and a localizer, they played each of the three games for five sessions. Each session lasted 7 minutes of gameplay, corresponding to 18,900 frames. Multiple episodes of the game could be played within a single session. An episode refers to the time from the start of the game until ‘game over’. Participants were allowed to take a short break between sessions.

In Breakout, the player controls a paddle at the bottom of the screen to bounce a ball and break bricks at the top. Hitting bricks scores points, while missing the ball results in losing one of five lives. In Space Invaders, the player controls a cannon to prevent aliens from invading. Destroying enemy ships earns points. Being hit or letting invaders reach the Earth results in the loss of one of three lives. In Enduro, the player steers a car in a racing game, earning points for successfully overtaking other cars and losing points if overtaken. Players must overtake a specified number of cars within a given time. Failing to do so results in ‘game over’. After reaching the required number of overtakes, no further points can be earned until the start of the next day.

A button box with four buttons arranged in a slightly curved row was used as controller. The first two buttons on the left were assigned to ‘fire’ and ‘brake.’ In Enduro, the ‘fire’ button controlled acceleration, in Space Invaders it fired the cannon, and in Breakout it launched a new ball. This button was operated by the left middle finger. The second button, used for braking, was only functional in Enduro and was controlled by the left index finger. The two buttons on the right controlled steering: the third button, operated by the right index finger, controlled ‘left,’ and the fourth button, operated by the right middle finger, controlled ‘right.’ Combinations of these actions were possible in Space Invaders and Enduro. In Breakout, we slowed down the paddle’s steering by applying the action taken by the participant only every second frame.

#### 2.2.3 Task implementation and data collection

The game instructions were implemented using the Tkinter library, while the Arcade Learning Environment (ALE; Bellemare et al., 2013), combined with the Python interface (bbitmaster, 2015), was used to present the games and log all relevant game data. This included motor responses, received rewards, the number of episodes played per session, and the presented screens. The relevant game information was stored as a time series with a temporal resolution of 45 Hz. The observed screens were saved as sequences of matrices representing grayscale pixel intensity values. The interface was modified and adapted to match the experimental design. The experiment was presented on a 1024 × 768 pixel screen (width × height), with states updated at a frequency of 45 Hz. Each state consisted of the maximum pixel values from the current and previous frames to reduce flickering. States were displayed in grayscale at a resolution of 84 × 84 pixels, consistent with the input used for the DQNs. Only the scoreboard was shown at a higher resolution to enhance readability.

### 2.3 fMRI data aquisition and preprocessing

#### 2.3.1 fMRI aquisition parameters

The dataset was acquired at the Neuroimaging Center of the TUD Dresden University of Technology. fMRI data were collected using a 3T MRI scanner (Siemens MAGNETOM Prisma) equipped with a 32-channel head coil. Initially, a field map was acquired to correct for echo-planar imaging (EPI) distortions, with repetition time (TR) = 749 ms, short echo time (TE) = 4.92 ms, long TE = 7.38 ms, flip angle = 56°, slice thickness = 2 mm. For whole-brain functional imaging, simultaneous multi-slice (SMS) acceleration with an SMS factor = 6 was used, with TR = 987 ms, TE = 32 ms, flip angle = 62°, FOV = 192 mm, 72 slices, and a voxel size = 2 mm isotropic. On the first day, a T1-weighted anatomical reference scan was collected with a TR = 1900 ms, TE = 2.26ms, inversion time (TI) = 900 ms, flip = 9°, and a spatial resolution = 0.5 mm × 0.5 mm × 1.0 mm.

#### 2.3.2 fMRI preprocessing pipeline

After data acquisition, the fMRI data were preprocessed using SPM12 running on MATLAB R2021b. First, slice-time correction was applied to the functional images, considering the SMS acquisition sequence. The images were then spatially realigned and unwarped using a voxel displacement map, calculated from the field map, to correct for distortions. The high-resolution structural image of the participant was co-registered to the mean functional image and segmented. The structural and functional images were normalized to the standard Montreal Neurological Institute (MNI) space, with a resampled voxel size of 2 mm isotropic. Spatial smoothing was applied using a Gaussian kernel with a full-width at half-maximum (FWHM) of 6 mm isotropic. In addition to the image realignment, a separate general linear model (GLM) was calculated for motion correction, using six motion parameters (three translations and three rotations) as regressors, with the preprocessed fMRI signal as the dependent variable (Friston et al., 1996). A high-pass filter with a cutoff of 128 seconds was applied as part of the GLM specification. This procedure accounted for motion-related signal fluctuations, and the resulting residuals were used as head motion-corrected fMRI data in our analyses. The correlation maps were interpolated using trilinear interpolation for visualization.

### 2.4 Analysis

#### 2.4.1 DQN-based encoding model analysis

To model voxel activities underlying S-R transformations, we used a DQN-based encoding model approach (Güclü & van Gerven, 2015). Our encoding model analysis consisted of two components: a DQN, which nonlinearly mapped the presented stimulus into a feature space, and a GLM, which mapped these DQN-generated features into the voxel space to predict neural activity. Gameplay screens generated by the subjects were processed through a DQN that had been specifically trained on the game. This resulted in a time series of activation values for each neuron across all layers of the network. Each time point in this series corresponded to the activation of a neuron in response to the current input presented to the model. To predict the fMRI time series, we fitted a voxel-specific regularized GLM, which mapped the generated stimulus features onto the fMRI data. The time series of the DQN’s activation values served as predictors, while the voxel’s response served as the dependent variable (see Figure 1). Depending on the object of investigation, the GLM predictors were selected either by including the activations of all neurons across all layers of the DQN (as in Sections 3.1 and 3.2) or layer-wise, using only the time series of neurons from a specific layer (as in Sections 3.2 and 3.3). We evaluated prediction accuracy using a 5-fold cross-validation procedure. The GLM was fitted to data from four out of five sessions.

**Figure 1:**
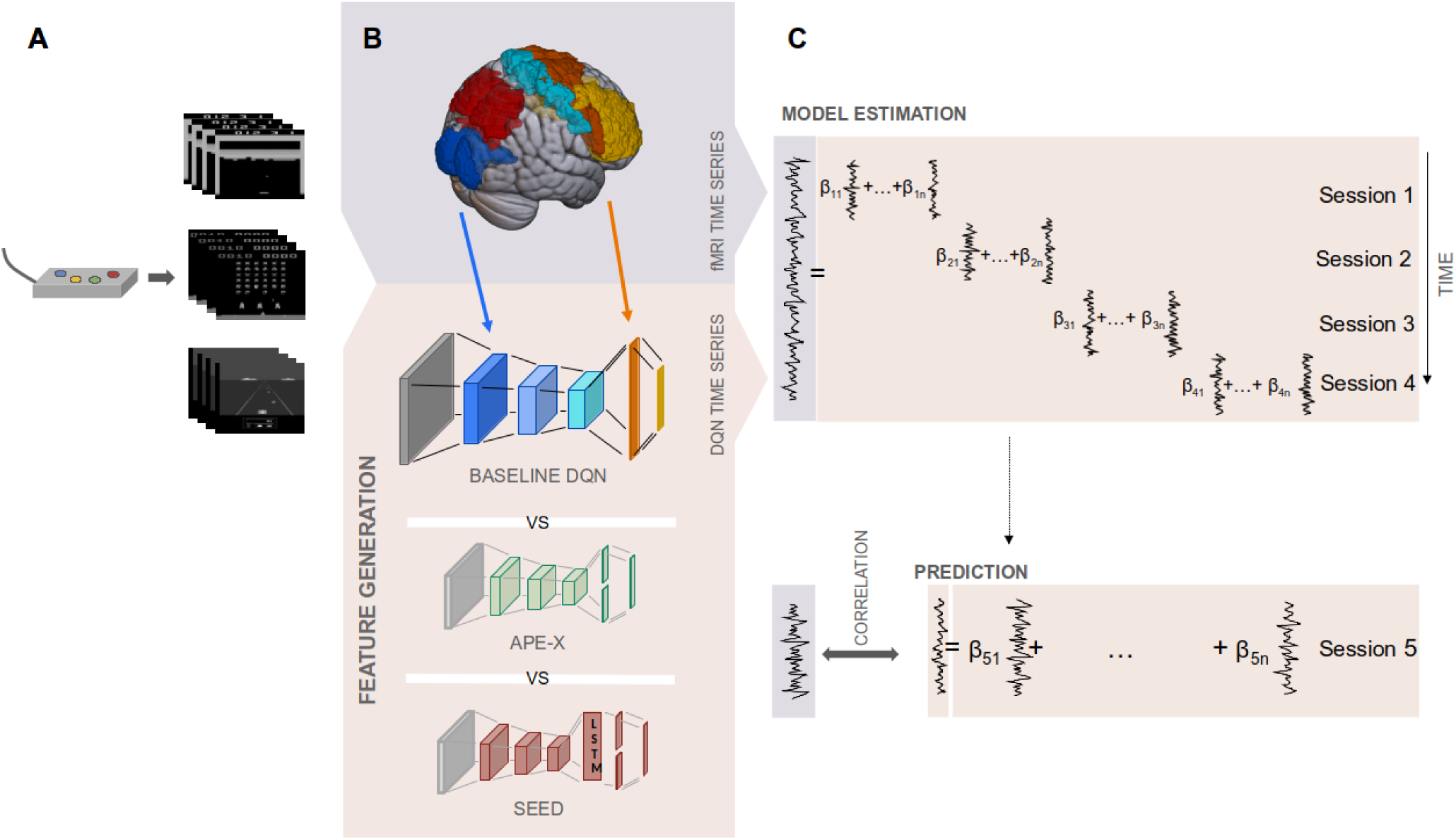
Analysis pipeline using a DQN-based encoding model. (A) Humans played arcade games inside an fMRI scanner using a four-button controller. (B) Video frames were processed through the DQNs, generating a time series of activation values for each neuron across all layers. Schematic representation of the color-coded hierarchical correspondence between DQN layers and brain regions (V1/V2 = blue, PPC = red, M1 = light blue, PMC = orange, LPFC = yellow), further analyzed in Section 3.3. In addition to the stimulus input, the DQN comprises three convolutional layers (layers 1-3), one fully connected layer (layer 4), and an output layer. In Ape-X and SEED, the fourth layer implements a dueling architecture. In SEED, an additional LSTM layer is inserted between the third convolutional layer and the fourth layer. (C) For each game, DQN, subject, and voxel, these stimulus features served as predictors in a GLM, fitted to the voxel’s fMRI time series using four of five sessions in a 5-fold cross-validation procedure. The fitted GLM was then used to predict the fMRI time series of the held-out session, with prediction accuracy assessed via Pearson correlation between actual and predicted fMRI time series.

Then we averaged the resulting beta weights and used them, together with the DQN-derived predictors from the held-out session, to predict the voxel’s time series in that session. The prediction performance was quantified by computing the Pearson correlation coefficient between the predicted and the actual fMRI time series. This procedure was repeated such that each session served once as the test set, while the remaining sessions were used for training. We compared the prediction accuracy of three encoding models, differing in the DQN component, using data from three games.

#### 2.4.2 DQN component of the encoding model analysis

DQNs are DNNs that combine deep learning with a model-free RL technique to solve complex tasks in environments with large and high-dimensional state spaces, such as arcade games (Mnih et al., 2015). The Q-learning algorithm is used to approximate the Q-value function, where the Q-value represents the expected discounted future reward for each (state, action)-pair based on the rewards obtained from executing an action in the given state (Watkins & Dayan, 1992). The network takes the state as input and outputs the Q-values for all possible actions and the action with the highest Q-value is then selected to maximize the reward and solve the task.

In our encoding model analysis, the DQNs served as non-linear, feature-generating mappings. We investigated three different DQNs, each representing a distinct stage of development and complexity. In our study, we used the baseline DQN as our baseline (Mnih et al., 2015). This architecture represents a basic implementation of DQNs. Ape-X and SEED (Espeholt et al., 2019; Horgan et al., 2018) build upon the baseline, incorporating advanced training procedures and architectural improvements to enhance performance and training efficiency. SEED, the most advanced of the three models, extends Ape-X further through additional optimizations. All three DQNs share a common structure, starting with three convolutional layers with rectified linear (ReLU) activations. The first convolutional layer contains 20 × 20 × 32 units, followed by a second layer with 9 × 9 × 64 units, and a third layer with 7 × 7 × 64 units. Each DQN includes an output layer whose neurons correspond to the possible actions. At each time step, the DQN received a stack of four consecutive video frames as input and information on ‘game over’ states. For detailed information on the preprocessing of input frames, we refer to Section 3.2.2 and Section 3.2.3 in (Haberland et al., 2025). Here, we provide only a brief overview.

**Table 1:** Overview of the three DQN architectures, emphasizing their similarities and differences.

| Baseline DQN | Ape-X | SEED |
| --- | --- | --- |
| **Input:** |  |  |
| Received a stack of four consecutive video frames and a terminal state flag as input. | Received a stack of four consecutive video frames and a terminal state flag as input. | Received a stack of four consecutive video frames, a terminal state flag, the achieved reward as input, and maintains past game states via an LSTM. |
| **Architecture:** |  |  |
| Vanilla feed-forward architecture: after convolutional layers (layers 1-3), a fully connected layer with 512 units is added (layer 4) before the output layer. | Dueling network architecture (Wang et al., 2016): after convolutional layers (layers 1-3), two separate streams with 512 units each estimated the state value and action advantages (layer 4), which are combined in the output layer. | Recurrent dueling architecture: after convolutional layers (layer 1-3) and a fully connected layer, an LSTM module (Hochreiter & Schmidhuber, 1997) with a single LSTM cell with 512 units is applied to capture temporal context for future decision-making, enabling the model to determine which information is relevant. This is followed by a dueling architecture comprising two separate streams of 512 units each (layer 4) and the output layer. |
| **Training methods:** |  |  |
| The gaming experiences were stored in a replay buffer (Lin, 1992), and random batches were sampled to train the network. | The training was distributed across multiple actors and a learner. Actors collected experiences in parallel environments, prioritized using prioritized replay (Schaul et al., 2016). Double Q-learning (van Hasselt et al., 2016) and multi-step bootstrap targets (Sutton, 1988) were applied. | The training was coordinated by a centralized learner responsible for inference, trajectory accumulation, and the computation and updating of model parameters, while actors executed actions in multiple environments and sent their observations back to the learner. It combined Q-learning with policy gradient methods and integrated double Q-learning, multi-step targets, prioritized replay, value function rescaling (Pohlen et al., 2018), and V-trace (Espeholt et al., 2018). Optimized task distribution, efficient resource utilization, and advanced parallelization strategies improved training stability and scalability. |
| **Training time:** |  |  |
| Three days of training and approximately 60 million environmental frames. | Eight days of training and approximately 800 million frames. | Due to limited computational resources, pre-trained network checkpoints from (Espeholt et al., 2021) were used, corresponding to approximately 40 billion frames. |
| **Gaming performance:** |  |  |
| Achieved human-level performance in Enduro. | Exceeded human performance in all three games. | Exceeded human performance in all three games. |
| **Code:** |  |  |
| The implementation follows the theoretical framework presented in (Mnih et al., 2015). | The code is based on the GitHub repository (neka-nat, 2020), with modifications for integration into our data analysis pipeline and the arcade environment. | Code and checkpoints were obtained from the GitHub repository (Espeholt et al., 2021). |

#### 2.4.3 GLM component of the encoding model analysis

After generating the stimulus features and preprocessing the fMRI data, we used voxel-specific regularized GLMs as linear encoding models. These models were fitted on training data to predict voxel activity in the test set across each DQN, game, and participant. The selection of predictors, whether layer-specific or derived from all layers, was determined by the research objectives. Only the time series of neurons that showed activation at least once during feature generation were included as predictors. We used activations from the convolutional layers following the application of the ReLU activation function. The number of predictors in the design matrix varied depending on the layer, game, DQN model, and subject. On average, the layers contributed approximately: ∼10,000 for the first, ∼5,000 for the second, ∼3,000 for the third, and ∼500 for the fourth layer. In SEED, the LSTM layer contributed additional ∼500 predictors prior to any regularization (see Section 2.4.2 for layer details). The design matrix, composed of these neuronal time series, was standardized along the temporal and feature dimensions by converting them into z-scores. To account for the hemodynamic delay, the regressors were convolved with a hemodynamic response function (HRF). To match the generated features with the fMRI sampling rate (TR frame rate), predictors were downsampled to the EPI frequency and a high-pass filter with a cutoff of 128 seconds was applied. Before solving the GLM, the predictors and the fMRI time series were z-scored again along the temporal dimension. Because the number of regressors exceeded the number of available data points (except in GLMs restricted to output layer neurons), it was necessary to regularize the GLM during the training phase of the 5-fold cross-validation, extending the standard GLM to a regularized GLM. In this context, L1-regularization proved most effective (Mohr & Ruge, 2021). Unlike unregularized or L2-regularized models, L1-regularization requires iterative solving. The model was solved using the MATLAB toolbox lasso gpu (Mohr & Ruge, 2021) and the regularization parameter was selected based on the mean cross-validation performance across all voxels and then held constant for all voxels. Voxel responses in the test set were predicted using a linear combination of the test-set features and weighted by the mean regression coefficients estimated from the training folds. The predicted voxel time series was then compared to the measured voxel time series using Pearson correlation. For each voxel, prediction accuracy was calculated as the average Pearson correlation across participants, games, and test sessions, separately for each DQN-based encoding model. We normalized the confidence intervals according to (Morey, 2008).

#### 2.4.4 Selection of regions of interest

To investigate brain activity along the dorsal stream, we defined a set of regions of interest (ROIs). The early visual ROI was based on the Jülich Brain Atlas and included regions V1 and V2. The posterior parietal cortex (PPC) was defined using the AAL2 atlas and included the precuneus and superior parietal lobule (SPL). The primary motor cortex (M1) was defined anatomically as Brodmann area 4 (BA 4), and the premotor cortex (PMC), including the supplementary motor area (SMA), corresponded to Brodmann area 6 (BA 6). The lateral prefrontal cortex (LPFC) included the middle frontal gyrus, inferior frontal gyrus pars triangularis, and pars opercularis, also based on the AAL2 atlas. The dorsal ROI encompassed the PPC, M1, and PMC. To enable comparison with the ventral visual stream, the ventral ROI included the inferior temporal cortex (IT) and middle temporal gyrus (MTG), again based on the AAL2 atlas. All ROI masks were created using the WFU PickAtlas toolbox in SPM12.

### 2.5 Software and hardware

The experiment and analysis were conducted on Ubuntu 20.04 as the operating system (see also Section 2.2.3). The experiment was run within a Docker container based on the python:3.9.7-buster image, which included Python version 3.9.7 and was built on Debian 10 (Buster). The DQNs were trained and used for feature extraction within Docker containers using the ALE framework (Bellemare et al., 2013) and its Python interface (bbitmaster, 2015). The baseline DQN and Ape-X were trained on Debian ‘Buster’ using Python 3.8.8 and PyTorch. SEED was trained with Python 3.6.9 and TensorFlow, running on Ubuntu 18.04.5 LTS. All analyses and estimations of the regularized linear models were performed in MATLAB 2019a. Our computational setup included an NVIDIA RTX A4000 GPU with 16 GB of memory and an Intel Xeon W-2245 CPU with 8 cores and 64 GB of RAM.

## 3 Results

To investigate S-R transformations, we analyzed fMRI data from *N* = 23 human participants while they played the arcade games Breakout, Space Invaders and Enduro. To predict voxel activity, we used a DQN-based encoding model approach (see Figure 1). First, we employed a DQN as a feature-generating mapping, processing the video screens seen by the subjects through the DQN to generate time series of activation values for each neuron in each layer. Second, we implemented a GLM as our encoding framework, using the generated stimulus features as predictors and the time series of voxel activations as the dependent variable. Within a 5-fold cross-validation procedure, we fitted these features to the measured fMRI time series during the training phase. We evaluated their prediction accuracy during the test phase by calculating the Pearson correlation *ρ* between the actual measured and the GLM-predicted fMRI time series for each voxel. To assess whether advancements in RL could improve the accuracy of modeling voxel activity, we compared the prediction accuracy of three DQN-based encoding models, each differing in their feature-generating component. We evaluated the predictive performance for each subject on three arcade games on which the DQNs were trained without any human data. To ensure generalizability beyond a single task, we averaged the results across all subjects and the three games.

### 3.1 DQN-based encoding models predict voxel activity of task-related S-R transformations across the dorsal stream

To localize brain regions involved in transforming time-continuous visual stimuli into motor responses, we applied our encoding models using the activations from all layers of the DQN as predictors in the GLM. All three encoding models yielded significant predictions of voxel activity along the dorsal visual stream (FWE-corrected, p*<* 0.05). All models were able to capture brain activity in the early visual cortex, progressing through higher-order visual areas, and extending into the PPC and further into motor-related regions. In the frontal cortex, activations covered the precentral gyrus and the SMA, including M1 and PMC. Activations also extended into the LPFC (see Figure 2 and see Supplementary Table S11-Table S13 for activation peaks).

**Figure 2:**
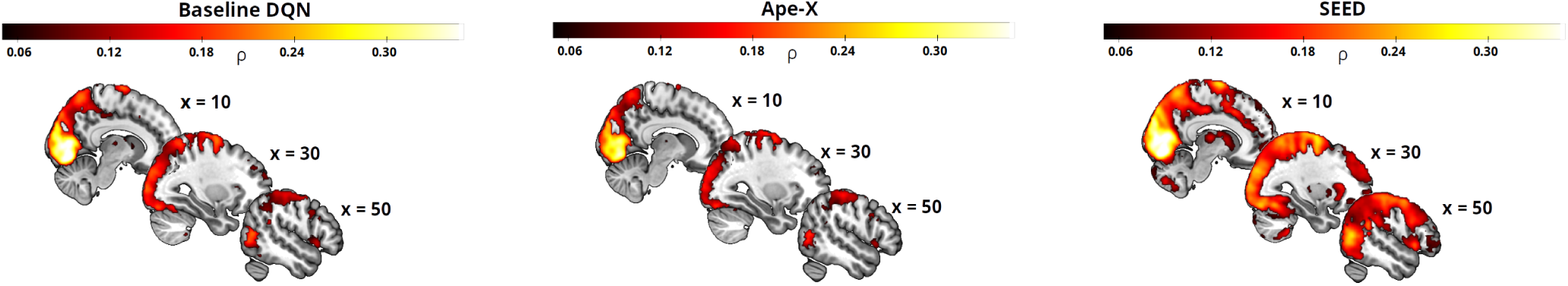
Prediction accuracy of the three encoding models. The plots show are whole-brain Pearson correlations between the actual and predicted fMRI voxel time series for the three encoding models, using feature maps from the baseline DQN (left), Ape-X (middle), and SEED (right). Neurons from all layers of the DQNs were used as predictors. High prediction accuracies were primarily observed in regions along the dorsal stream. Only statistically significant voxels are shown (FWE-corrected, p *<* 0.05).

All three encoding models predicted voxel activity significantly higher in the dorsal stream compared to the ventral stream (two-sample t-test, p *<* 0.001, see Figure 3), thereby replicating the well-established hypothesis that visuomotor tasks are predominantly associated with the dorsal stream rather than the ventral stream (Goodale, 2011; Kravitz et al., 2011). These results provided initial validation that DQN-based encoding models enable the modeling of time-continuous S-R transformations using high-dimensional stimuli in biologically plausible brain regions. Moreover, they offered preliminary evidence for similarities between the internal feature representations of the DQNs and neural activity in the human brain, motivating a more detailed layer-wise analysis to characterize the step-wise S-R transformation process.

**Figure 3:**
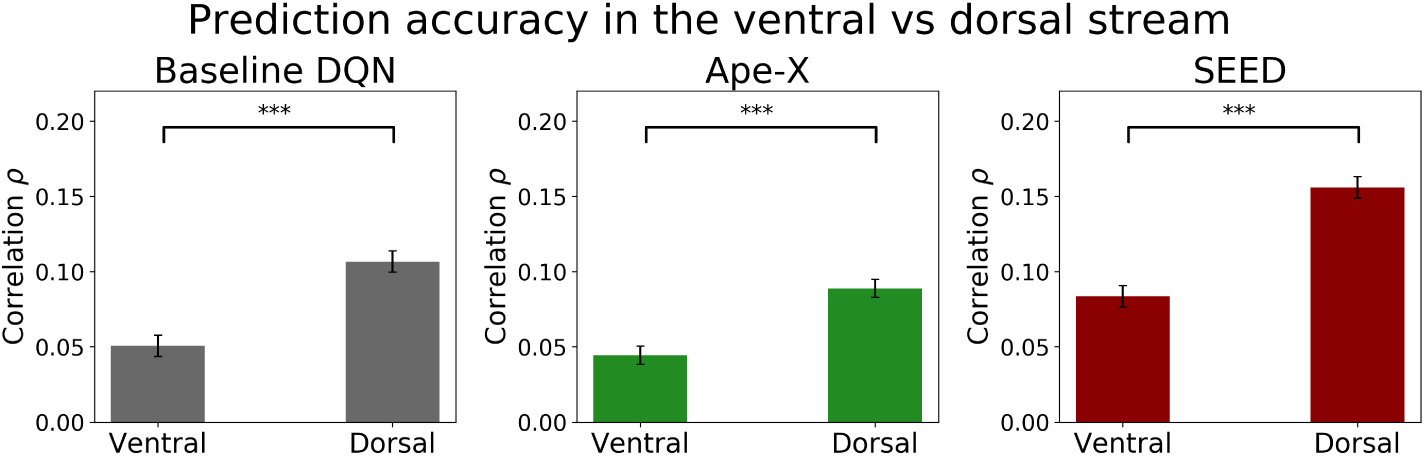
Bar plots show the voxel-wise Pearson correlation in ventral and dorsal regions, averaged across all voxels within the ventral (MTG, IT) and the dorsal (precuneus, SPL, M1, and PMC) stream. The encoding models used features from all layers of the baseline DQN (left), Ape-X (middle), and SEED (right). Error bars represent the normalized 95% confidence interval of the mean correlation across subjects. Prediction accuracies were significantly higher in the dorsal visual stream compared to the ventral visual stream (two-sample t-test, p *<* 0.001; significance denoted by ‘***’).

### 3.2 Features from advanced DQNs enable more accurate predictions of voxel activity

We demonstrated that the feature representations of all three DQNs were well-suited for predicting neural activity along the dorsal visual stream during gameplay. By comparing the encoding performance of the three DQNs and their strengths and limitations, we assessed the extent to which recent advances in machine learning contribute to improvements in neuroscientific modeling, potentially offering insights into the representational structure and computational principles of the human brain. We analyzed the differences in prediction accuracy across the three encoding models and subsequently investigated the origins of these differences. Our analysis revealed that SEED was the most effective feature-generating mapping, significantly outperforming the baseline DQN and Ape-X in terms of prediction accuracy along the dorsal stream (FWE-corrected, p *<* 0.05, see Figure 4 (A) and see Supplementary Table S14-Table S16 for activation peaks). Differences in predictions between the DQNs were observed in all investigated ROIs (repeated-measures ANOVA with the factors *ROI* and *DQN*, Greenhouse-Geisser corrected, revealed significantly main effects of ROI: *F*_ROI_(2.23) = 129.34, 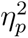 = 0.86, and DQN: *F*_DQN_(1.42) = 156.62, 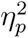 = 0.88, and a significant *ROI* × *DQN* interaction: *F*_ROI×DQN_(4.78) = 17.87, 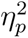 = 0.46, all p *<* 0.001; post hoc comparisons indicated that, within each *ROI*, SEED achieved significantly higher prediction accuracy than the baseline DQN and Ape-X, Bonferroni-corrected, p *<* 0.001, see Figure 4 (B)). These differences in prediction accuracy between the features of SEED and the other DQNs were particularly pronounced in higher-level visual and control regions, such as the PPC and PMC and less pronounced in lower-level areas, including V1/V2 and M1 (custom contrasts, Bonferroni-corrected, all p*<* 0.05, except PPC vs. V1/V2, after Bonferroni-correction *p* = 0.092, see Supplementary Table S4 and Table S5). No considerable differences were observed between the features of Ape-X and the baseline DQN, with only a few voxels reaching statistical significance, suggesting that Ape-X and the baseline DQN produce similar feature spaces in terms of prediction accuracy (FWE-corrected, p *<* 0.05, see Figure 4 (A) left). While all three DQNs were able to generate features that predict voxel activity in lower-level regions, higher-level regions appear to require more abstract or complex features, captured only by the most advanced model, SEED.

**Figure 4:**
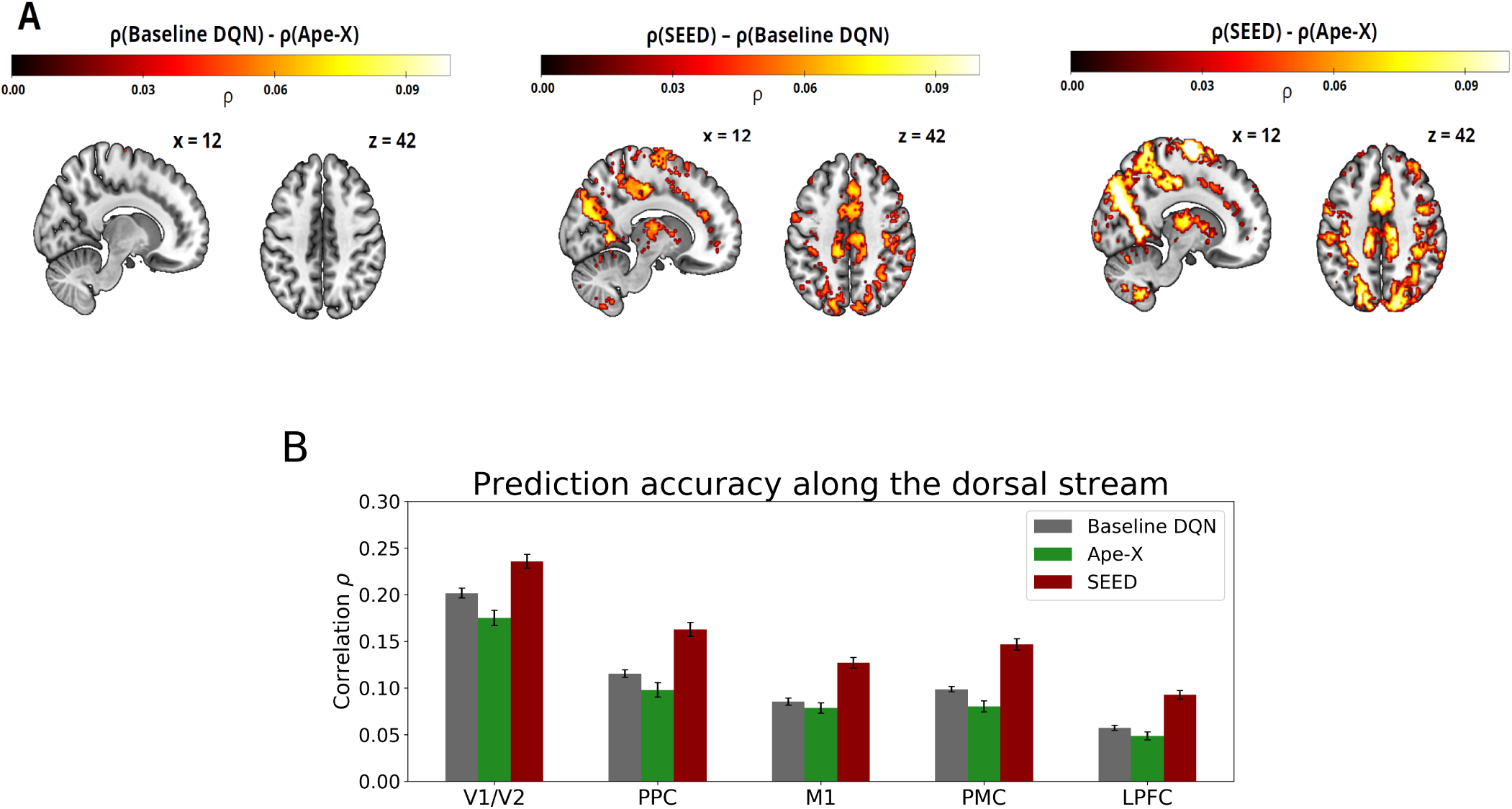
(A) Differences in prediction accuracy between the three encoding models. Intensity values reflect significant differences in voxel-wise Pearson correlations, highlighting regions where the feature representations of one DQN outperform those of another in predicting voxel activity (FWE-corrected, p *<* 0.05). No considerable differences were observed between Ape-X and the baseline DQN (left). SEED outperformed the baseline DQN (middle) and Ape-X (right). Features from all network layers were used as predictors in the encoding models. Reverse subtractions revealed no voxels with significant positive differences (FWE-corrected, p *<* 0.05). SEED generated the most effective feature representations for predicting voxel activity, showing the greatest improvements in voxels within the PPC and PMC regions. (B) Prediction accuracy of the three encoding models across ROIs. Bar plots show mean voxel-wise Pearson correlations within each ROI. Encoding models were based on features from the baseline DQN (gray), Ape-X (green), and SEED (red). The ROIs encompass V1/V2, PPC (precuneus, SPL), M1 (BA 4), PMC (BA 6), and LPFC (middle frontal gyrus, the inferior frontal gyrus pars triangularis, and pars opercularis). Error bars represent the normalized 95% confidence interval of the mean correlation across subjects. The improved prediction based on features generated by SEED was particularly evident in higher-level visual and control regions.

After demonstrating that advances in RL can be leveraged to improve modeling in neuroscientific research, we performed a layer-wise breakdown and comparison of the DQNs’ encoding capabilities. This allowed us to identify the hidden network layers that contributed most substantially to the observed differences and to gain insight into the functional role of each computational stage. We addressed this by replacing the GLM, which included neural activity from all DQN layers as predictors, with voxel-specific GLMs incorporating layer-specific predictors (see Figure 5). All three DQNs shared an identical architecture in their initial layers, consisting of three convolutional layers. As the second convolutional layer exhibited characteristics highly similar to those of the first and third, we focused our analysis on the first and third layers to maintain clarity. This structural similarity was reflected in the layer-wise prediction performance: although significant differences between the DQNs emerged already in the earlier layers (repeated-measures ANOVA for the first and third layer with the factors *DQN*, *Layer* and *ROI*, Greenhouse-Geisser corrected, revealed a significantly main effect of DQN: *F*_DQN_(1.91) = 10.32, 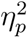 = 0.33, and a significant *Layer* × *DQN* interaction: *F*_Layer×DQN_(2.00) = 11.68, 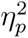 = 0.36, all p *<* 0.001), the magnitude of these differences remained small across the first and third layer in all examined ROIs. In contrast to the first three layers, the fourth layer showed significantly larger differences between the DQNs in prediction accuracy across the ROIs (custom contrast of the *DQN* × *Layer* interaction, p*<* 0.001, see Supplementary Table S6 and Table S7). This suggests that the early convolutional layers of the DQNs, despite differences in training regimes and architectural complexity, encode similar feature spaces, with the differences shown in Figure 4 being primarily attributable to SEED’s fourth layer, the layer in which architectural divergence becomes prominent. Differences in prediction accuracy between features generated by SEED’s fourth layer, which follows the LSTM, and those from the fourth layers of the baseline DQN and Ape-X were more pronounced in higher-level control regions, including the PPC and PMC, than in lower-level areas such as V1/V2 (custom contrasts, Bonferroni-corrected, p *<* 0.01, see Supplementary Table S8 and Table S9, and see Supplementary Figure S7 and Figure S8). This pattern suggests a potential functional relevance of the LSTM component in capturing representations related to higher-level cognitive control processes.

**Figure 5:**
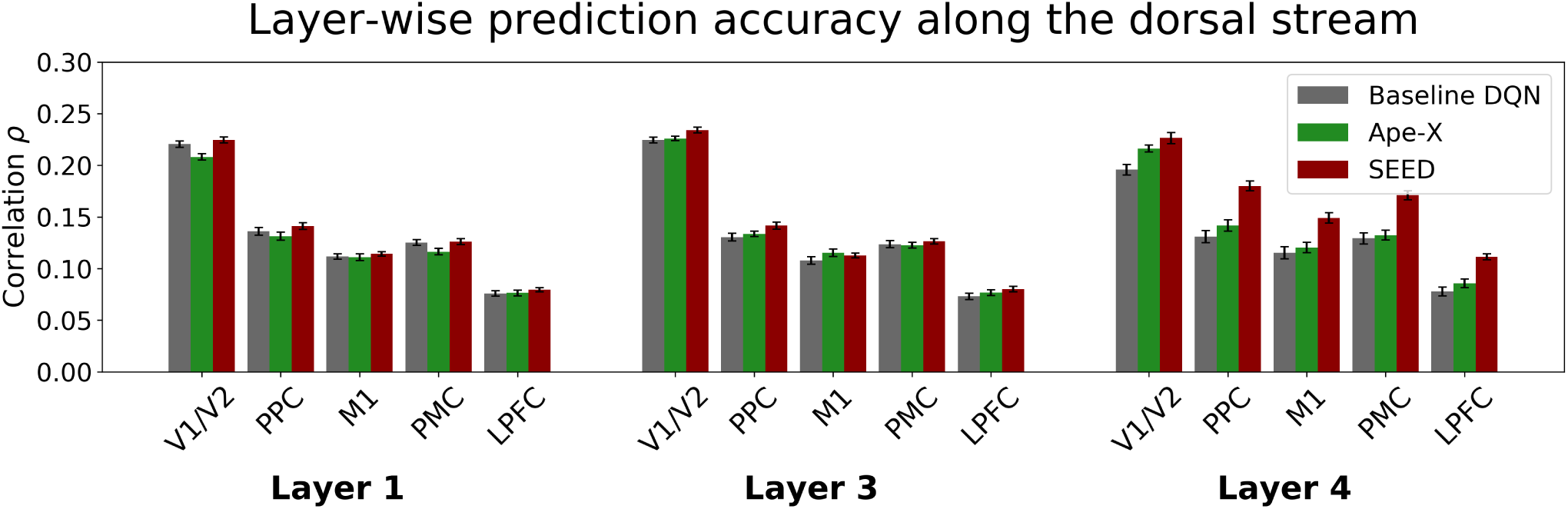
Layer-wise comparison of prediction accuracy. The bar plot shows the Pearson correlation between actual and predicted time series for individual layers for voxels within V1/V2, PPC, M1, PMC, and LPFC. The analysis included DQN neurons from the first, third, and fourth layers, using encoding models based on features from the baseline DQN (gray), Ape-X (green), and SEED (red). Error bars represent the normalized 95% confidence interval of the mean correlation across subjects. Breaking down predictive performance by layer enabled a direct comparison of representational alignment between models and brain regions, offering insights into which stages of computation most closely resemble neural processing. Architectural differences among the DQNs, particularly in the fourth layer, were reflected in prediction accuracy. Notably, SEED’s fourth layer generated features that better explain voxel activity, particularly in higher-level control regions, than those of the baseline DQN and Ape-X.

### 3.3 Advanced DQNs capture brain-like hierarchical processing stages

Since SEED has proven to be a suitable feature-generating mapping for predicting voxel activity along the dorsal stream, we focused our further analysis on SEED to investigate whether it captures the different stages of human visuomotor processing. Understanding how SEED’s internal representations align with the brain’s processing hierarchy can deepen our understanding of neural coding and may help characterize the functional organization of neural computations in this stream from a computational standpoint. To assess whether the hierarchical structure of SEED aligns with the organization of the dorsal visual stream, we compared the voxel-wise prediction accuracy by assigning each voxel to the SEED layer that yielded the highest layer-wise prediction accuracy (see Figure 6). This voxel-wise mapping revealed that most voxels best predicted by the three early convolutional layers were located in V1/V2 and M1. In the low-level visual areas V1/V2, the ratio of early (first, second, and third) to late (LSTM and fourth) layers was significantly higher than in the higher-level ROIs, whereas the ratio decreased in regions such as the PPC, PMC, and LPFC (pairwise t-tests for the factor *ROI* on the ratio of early layers to late layers, after applying the log-ratio transformation described in (Greenacre, 2021) to account for compositional data, p*<* 0.001, Bonferroni-corrected, see Supplementary Table S10). These results suggest that SEED captures brain-like hierarchical representations, with its layer-wise representations reflecting the cortical progression from early visual to higher-order visuomotor areas along the dorsal visual stream (see also Supplementary Figure S4).

**Figure 6:**
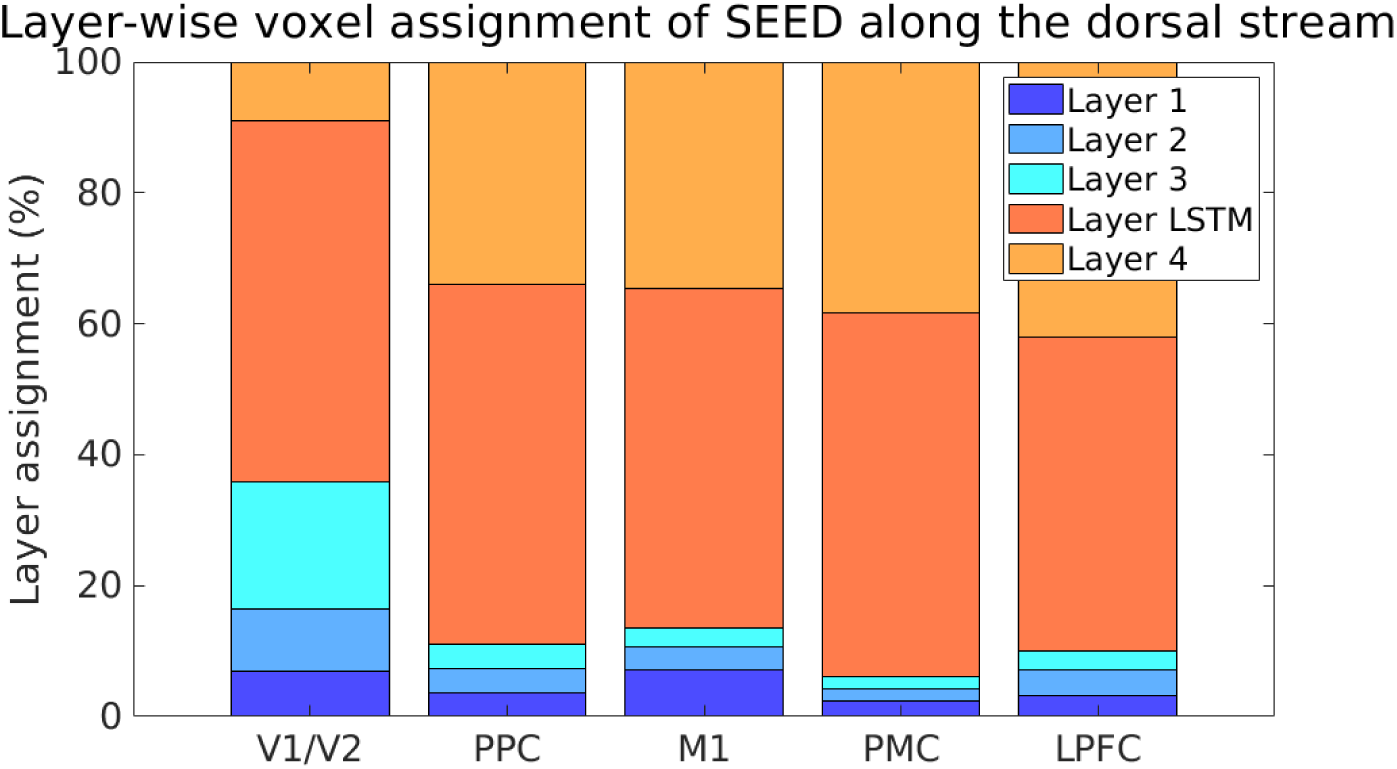
Voxel assignment to the layer of maximal prediction accuracy in SEED. To identify which hidden layers of SEED provided the most predictive features for different brain regions, each significant voxel (FWE-corrected, p *<* 0.05) was assigned to the hidden layer yielding the highest voxel-wise and layer-wise correlation between predicted and actual activity. The proportion of each layer in the bars (per ROI) reflects the percentage of voxels within that ROI assigned to the corresponding layer. This analysis was conducted for V1/V2, PPC, M1, PMC, and LPFC, following the approach in (Güclü & van Gerven, 2017a; Güclü & van Gerven, 2015). In V1/V2 and M1, the proportion of voxels predicted by the first convolutional layers was larger than in the other ROIs. In higher-level areas, such as PPC, PMC, and LPFC, the proportion of voxels associated with early layers progressively decreased, while the predictive contribution of the later layers increased. A strict one-to-one mapping between network layers and brain regions is unlikely because of the dependent nature of the network, where the layer’s output serves as input to the next.

## 4 Discussion

Understanding how the human brain continuously transforms complex, high-dimensional sensory inputs into motor responses within a dynamically changing environment is crucial for explaining everyday behavior. To gain a more detailed characterization of the neural representations underlying individual S-R processing stages, we analyzed human gameplay behavior during arcade games performed inside an fMRI scanner, using a DQN-based encoding model. We demonstrated similarities in internal representational feature spaces between artificial and biological neural networks. By comparing three DQNs with varying architectures and training methods, we showed that recent advances in machine learning can be leveraged to model aspects of S-R processing. We also found analogies to hierarchical information processing in the brain.

While encoding models have been widely used to model neural brain activity, the rapid evolution of deep learning methods has opened new possibilities for neuroscience. The integration of DNNs in encoding models enabled the characterization of the visual cortex using hierarchically organized object-recognition CNNs (Cichy et al., 2016; Yamins & Dicarlo, 2016).

Despite methodological advances, capturing task-relevant brain activity in naturalistic, interactive environments remains challenging. Video games offer a promising framework to address this issue: they are time-continuous, require active engagement, and involve a wide range of cognitive processes rather than isolated functions (Bellec & Boyle, 2019). Playing arcade games may engage more than simple S-R transformations. High-level cognitive processes may be involved, and behavior may appear goal-directed and anticipatory while playing these arcade games. To increase the likelihood of observing more automated processing, participants practiced the games prior to scanning to reach a stable performance level (Mohr et al., 2016). Atari games are also widely regarded as effective testing environments for RL. Therefore, we leveraged a DQN-based encoding model applied to Atari gameplay. Our findings support previous evidence that DQNs, when used as feature-generating mappings, enable the investigation of S-R transformations along the dorsal stream in time-continuous and complex environments, and that our results are consistent with those reported in (Cross et al., 2021).

The video game paradigm has received limited attention in the literature so far (Cross et al., 2021; Kemtur, 2023; Paugram, 2024; Tomov et al., 2023). These studies have shown that RL-trained models not only acquire task-solving skills but also develop effective encoding capabilities. Neither the architecture alone (see Supplementary Figure S9) nor a principal component analysis (PCA) of the raw pixel input is sufficient to predict brain activity (Cross et al., 2021). Instead, the training objective plays an important role in shaping model performance. When controlling for data quantity and type, RL outperforms imitation learning, which models human gameplay (Kemtur, 2023), and vision models in encoding brain activity during video gameplay (Paugram, 2024).

In this study, we focused on three different DQNs that shared the same training objective, maximizing reward through RL, but differed in training methods and architectural complexity. By comparing these models, it revealed their respective strengths and weaknesses as feature-generating mappings and identified key factors contributing to successful neural encoding. This, in turn, improves the ability to model neural mechanisms, which is essential for advancing the understanding of brain function (Cichy et al., 2016; Kriegeskorte, 2015).

When comparing the predictive performance of the features across the three DQNs, we found that the features from SEED, the most advanced of the three, outperformed those from the other DQNs. Differences emerged particularly in ROIs associated with higher-level cognitive control functions, such as the PPC and the PMC (Brovelli et al., 2015; Bunge et al., 2002; Podzebenko et al., 2002). In contrast, differences in prediction accuracy across the models were minimal in low-level areas, including V1/V2 and M1. This suggests that all three models are capable of representing low-level sensory features effectively. A layer-wise analysis revealed that the first three convolutional layers of the DQNs generated features yielding comparable prediction accuracies across ROIs. This confirms the assumption that differences in prediction performance do not arise from differences in the representation of low-level sensory stimulus features, but only become relevant at higher levels of abstraction (Güclü & van Gerven, 2015). In contrast, substantial differences emerged in the fourth layer, where SEED outperformed the baseline DQN and Ape-X, which both showed limited encoding performance in higher-level visual and control regions (see also Supplementary Figure S7 and Figure S8). These findings indicate that SEED’s higher layers develop more task-specific and architecture-sensitive representations, which are crucial for modeling brain activity in regions associated with higher-level cognitive control. This advantage is likely attributable to SEED’s architectural enhancements, specifically, the incorporation of an LSTM module, whose output is the input of the fourth layer, and the dueling network architecture.

LSTMs are RNNs that take temporal dependencies in data into account (Hochreiter & Schmidhuber, 1997). They filter task-relevant content using a system of gates, similar to mechanisms hypothesized for human working memory (Chatham & Badre, 2015; Frank et al., 2001). As a result, LSTMs are increasingly used as models to investigate working memory (Goldstein et al., 2019) and have been shown to capture functional activity of the visual working memory in the brain in N-back tasks (Sainath, 2020). Behavioral evidence suggests that incorporating an LSTM module is an important advancement in modeling human gameplay (Haberland et al., 2025). Our study suggests that temporally integrated representations of the LSTM improve the model’s sensitivity to temporal dynamics, improving its ability to capture brain-related features in higher-level regions, extending up to the LPFC (see also Supplementary Figure S7 and Figure S8). These areas are typically associated with cognitive control functions such as working memory maintenance and decision making (Smith & Jonides, 1999; Wager & Smith, 2003). However, compared to other regions, the correspondence between DQN-generated features and activity in the dorsolateral prefrontal cortex (dlPFC), a region typically associated with more abstract forms of cognitive control and the manipulation of working memory contents (D’Esposito et al., 1999), was less pronounced. This could reflect the relatively low demands on complex memory operations in Atari gameplay, which is largely stimulus-driven, or it may indicate limitations in the ability to capture such high-level cognitive functions with our models. Nonetheless, the observed patterns suggest that the LSTM layer may encode brain-relevant features of visual stimuli that align with the functional roles of these regions.

However, comparisons based on encoding model performance must account for potential confounding factors. Beyond model architecture, encoding performance is also substantially influenced by DQNs’ task performance (Schrimpf et al., 2018), their training dataset, and the amount of training (Paugram, 2024). In our study, such differences in training of the DQNs could therefore act as potential confounds. Consequently, the observed effects may, at least in part, reflect differences in training experience in addition to architectural characteristics. However, in our analysis, the fitting of the encoding components was always performed in the same way and on the same fMRI data across all three DQNs.

The layer-wise analysis across different DQNs not only highlights the role of recurrent structures but also reveals additional insights through the functional hierarchical correspondence between Seed’s network layers and the stages of S-R transformations along the dorsal stream (see also Supplementary Figure S3 and Figure S4). Early layers showed stronger alignment with the early visual cortex and, to a lesser extent, with M1, whereas higher-order regions such as the PPC and PMC aligned more closely with later layers (see Figure 6). This suggests that different brain regions are best characterized by distinct underlying feature spaces of varying complexity and it highlights the role of multiple stages of nonlinear feature transformations in capturing higher-order cognitive processes (Bengio, 2009; Creem-Regehr, 2009; Güclü & van Gerven, 2017a). Previous studies on video gameplay using DQNs without recurrent components found no evidence of a hierarchical gradient (Cross et al., 2021; Paugram, 2024). However, an imitation learning model that incorporated recurrent components showed hierarchical organization in video game tasks (Kemtur, 2023), supporting the idea that recurrence enables temporal dynamics to be captured and may be critical for hierarchical gradients. This functional correspondence between artificial and biological hierarchies provides a promising basis for exploring how different brain regions contribute to visuomotor transformations and how these computations are internally represented. At the same time, such insights may guide the development of more human-like DNNs with improved generalization capabilities, which in turn could serve as better computational models of brain function (Yamins & Dicarlo, 2016). Similar hierarchical correspondences have been observed in previous work on object recognition along the ventral (Eickenberg et al., 2017; Güclü & van Gerven, 2015; Nonaka et al., 2021) and dorsal (Cichy et al., 2016; Güclü & van Gerven, 2017a) visual streams. In our study, the convolutional layers did not show a hierarchy within the visual cortex, potentially reflecting discrepancies in task demands or architectural constraints (see Supplementary Figure S5). Prior research has also suggested that high-performing object recognition DNNs are not necessarily more brain-like (Nonaka et al., 2021; Schrimpf et al., 2018). Among the models we tested, SEED achieved the highest task and encoding model performance and showed the most pronounced representational gradient (see Supplementary Figure S3). This raises the question of whether a similar trade-off exists in more advanced DNNs trained on S-R transformation tasks, specifically, whether increasing task performance might come at the cost of human-like encoding.

Although the DQNs captured suitable representational spaces, they generated a large number of features in each layer, resulting in a relatively high number of GLM regressors compared to the limited number of data points, which made the models susceptible to the curse of dimensionality (Zahavy et al., 2016). To address this issue and make the estimation of the encoding models feasible, L1-regularization was applied (Ayinde & Inanc, 2019b; Jia et al., 2022), which has been shown to be the most effective approach for fitting a regularized GLM (Mohr & Ruge, 2021). Regularization substantially reduced multicollinearity among the GLM regressors, although some correlations within and across layers remained (see Supplementary Table S1-Table S3), which can influence GLM parameter estimates (Monti, 2011). Including features from all layers could therefore increase model complexity without necessarily improving predictive performance over layer-wise models (see Supplementary Figure S10). However, excessively strong regularization removed even important information, leading to a lower encoding performance. Reducing feature redundancy during DQN training could thus be a useful strategy, and several methods exist to quantify and mitigate this issue (Ayinde & Inanc, 2019a; Jia et al., 2022).

While these linear encoding models are data-efficient, easy to implement, and provide interpretable mappings between feature and brain activity spaces, further improvements in encoding could be achieved using nonlinear approaches such as recurrent networks (Güclü & van Gerven, 2017b; C. Zhang et al., 2019).

Our results contribute to the broader debate about whether DNNs are suitable models of cognitive processes, emphasizing the importance of critically evaluating their limitations while recognizing their potential. Critics argue that DNNs provide limited explanatory power for cognitive processes, as their internal processes are often opaque and difficult to interpret (Cichy & Kaiser, 2019). Furthermore, the development of DNNs is typically driven by engineering objectives rather than neurobiological theories, which limits their relevance as models of brain function (Kriegeskorte, 2015; Lake et al., 2017; Ma & Peters, 2020; Marblestone et al., 2016; Yamins & Dicarlo, 2016). This also contributes to their limited ability to generalize across tasks and also in capturing neural processes in new situations (Paugram, 2024). However, our analysis showed that DQN-based encoding models were able to predict neural activity and human motor responses with high accuracy, suggesting that these models capture cognitive processes that extend from neural representations to behaviorally relevant mechanisms (see Supplementary Figure S11) (Haberland et al., 2025; Mohr et al., 2019). However, DQNs that exhibit more human-like behavior do not automatically translate into better neural models at the individual level (see Supplementary Figure S6). This suggests that, while a DQN may capture aspects of behaviorally relevant processing, caution is important when interpreting these results, since explaining variance does not imply capturing the complexity of the brain’s functional mechanisms or underlying biological processes (van Gerven, 2017), a distinction that is therefore crucial (Shmueli, 2011). Against this critical background, however, they remain the most practical and general class of models, which are capable of learning to map high-dimensional input to appropriate motor responses, with initial evidence suggesting similarities in their representational spaces with the human brain. Their pragmatic value in cognitive neuroscience lies in their ability to scale to time-continuous, ecologically valid tasks that are often inaccessible to traditional analytical models. We were able to show that findings from highly controlled tasks also hold in more naturalistic visuomotor settings. Importantly, this is a highly exploratory approach that should be viewed as a valuable complement, not a replacement, for the traditional discrete and trial-based experimental paradigms. We argue that DNNs are still valuable scientific tools. A model should not only be interpretable but must also achieve sufficient predictive performance to justify making explanatory claims (Cichy & Kaiser, 2019).

Future developments in DNNs might benefit from integrating alternative assumptions, biologically inspired constraints, and objective functions that represent various theories of brain function. This may help align artificial dynamics more closely with neural processes (Bellec & Boyle, 2019; Federer et al., 2020). Additionally, advancements like multi-task learning (Parisotto et al., 2015) and transformer models (Vaswani et al., 2017) provide promising directions for further development. With ongoing improvements in interpretability, DNNs have the potential to serve as useful tools for testing hypotheses (Yamins & Dicarlo, 2016) and designing optimal stimuli (Bashivan et al., 2019). Analytical methods like representational similarity analysis (RSA) and neuroimaging tools like EEG can help to improve the understanding of how model activity is aligned with brain function in individual state-spaces. Analyzing specific game contexts can reveal latent subcomponents of behavior and reveal context-dependent S-R mappings. While our primary goal was to model S-R processing using brain-inspired artificial networks, our findings may also contribute to the development of more cognitively informed and efficient machine learning models.

Although challenges remain, our results provide additional proof of concept for a promising approach to investigating the neural representations underlying S-R transformations in time-continuous visuomotor tasks. This approach complements traditional trial-based paradigms by bridging controlled experimental research with real-world cognition. Our results demonstrate that cognitive neuroscience can benefit from advances in deep learning and encourage a broader methodological exchange between neuroscience and machine learning.

## Data and code availability

All behavioral data, the recorded screen observed by the subjects, as well as the results of our analysis on single-subject and group-analysis level are publicly available at https://osf.io/9cwq4/ (DOI: 10.17605/OSF.IO/9CWQ4).

The experimental tasks, the DQN code used to generate the features, and all analysis scripts are accessible via the public GitHub repository at https://github.com/SHaberland15/Arcade_DQN_Research_fMRI. A permanent DOI for the repository was created via Zenodo: 10.5281/zenodo.17085737.

## Author contributions

Sabine Haberland: methodology, software, validation, formal analysis, investigation, data curation, writing - original draft, and visualization. Holger Frimmel: conceptualization, methodology, software, writing - review and editing, and funding acquisition. Hannes Ruge: conceptualization, validation, formal analysis, resources, writing - review and editing, supervision, and project administration.

## Supplementary materials

**Figure S1.**
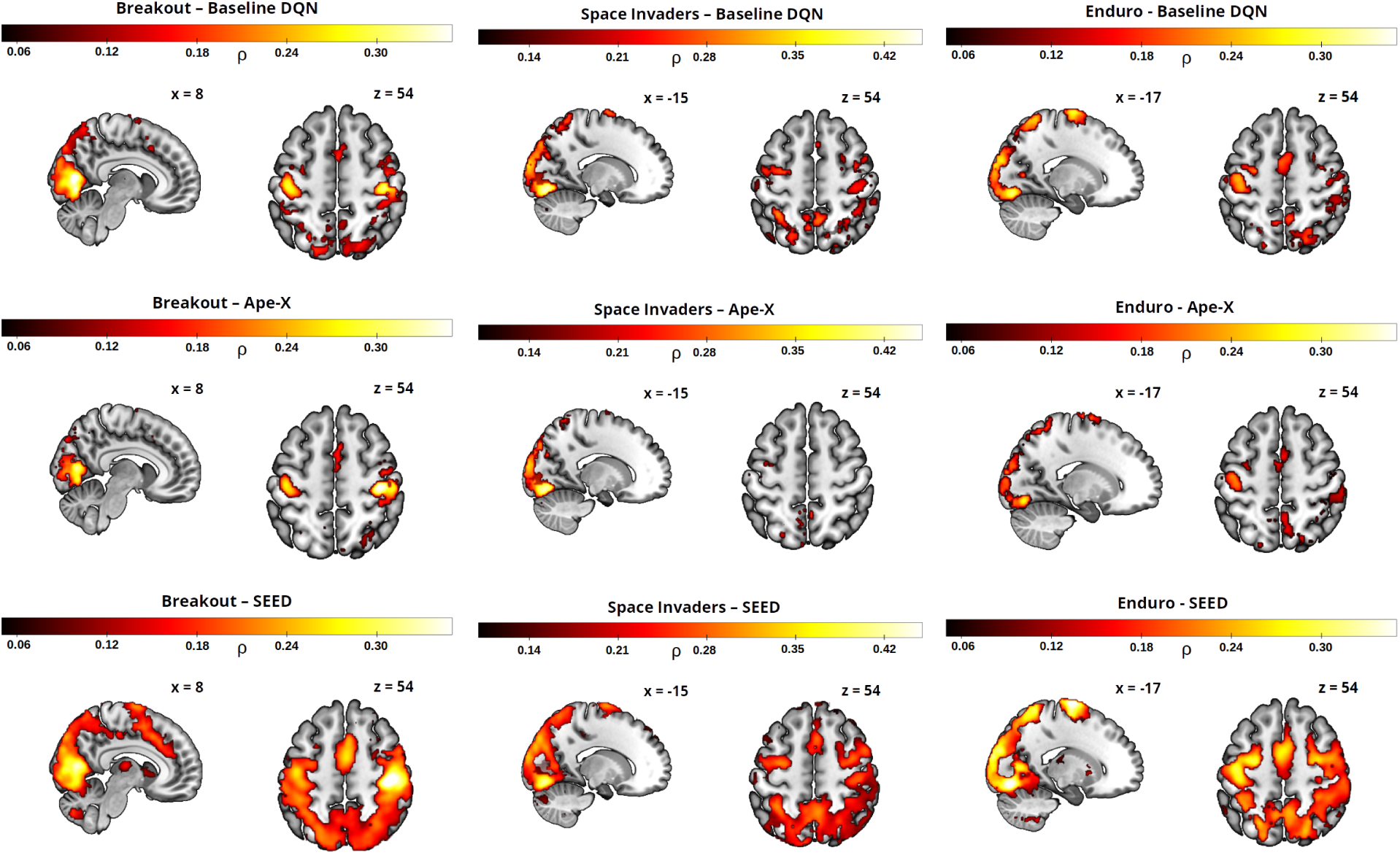
Prediction accuracy of the three encoding models across the three games. Pearson correlation between the actual and predicted fMRI voxel time series for the three encoding models using features from the baseline DQN (top), Ape-X (middle), and SEED (bottom). Neurons from all layers were used as predictors for each of the three games: Breakout (left), Space Invaders (middle), and Enduro (right). Only statistically significant voxels are shown (FWE-corrected, p *<* 0.05). While Figure 2 shows the mean prediction accuracy across all games, this figure shows the results separately for each game.

**Figure S2.**
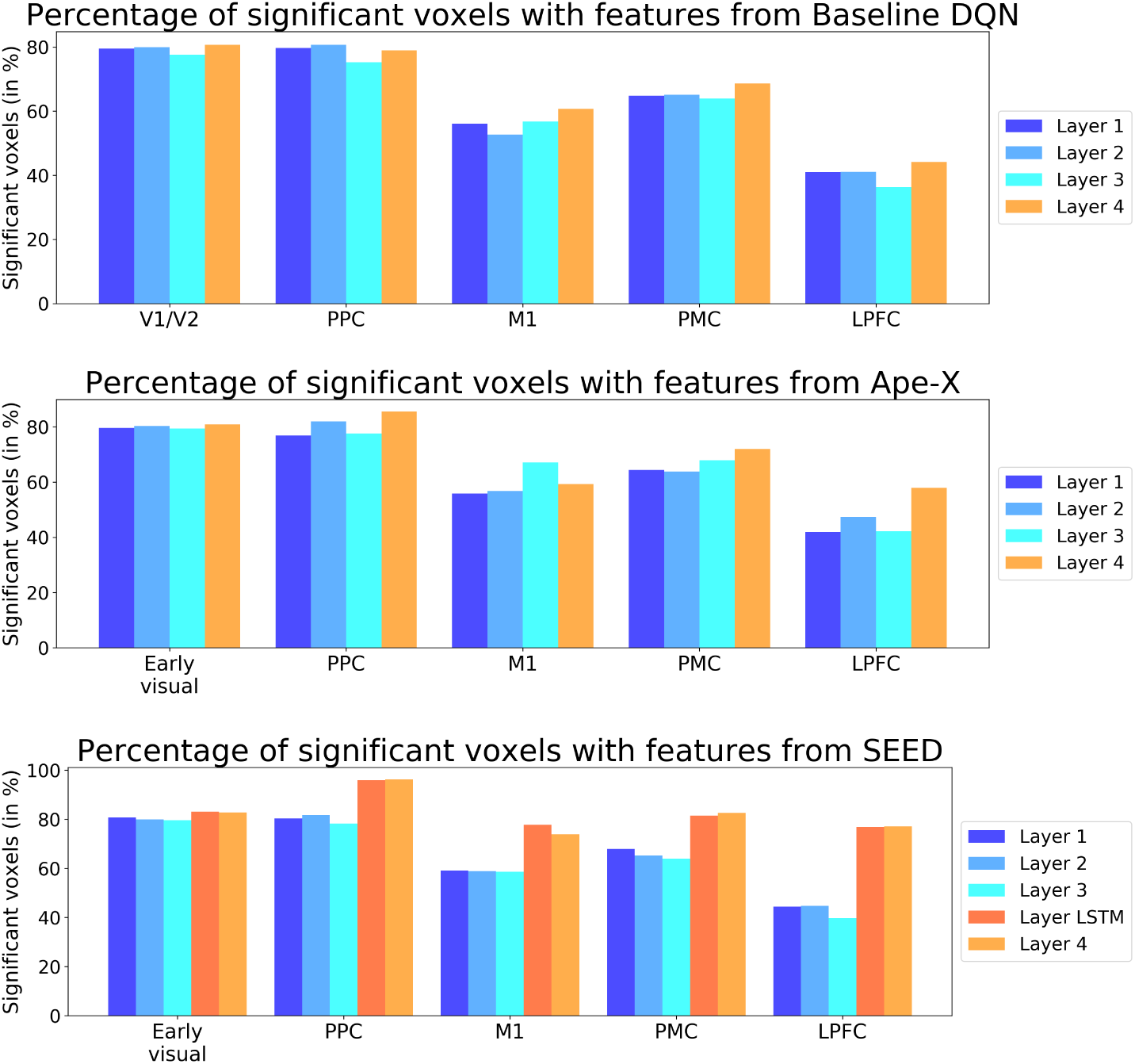
ROI-based analysis of voxels with significant prediction accuracy. The bar plots show the proportion of significantly predicted voxels within each ROI relative to the total number of voxels in that ROI. The analysis was conducted on DQN neurons from the first, third, and fourth layers, using encoding models based on features from the baseline DQN (left), Ape-X (middle), and SEED (right) (FWE-corrected, p *<* 0.05). The evaluated ROIs included V1/V2, PPC, M1, PMC, and LPFC.

**Figure S3.**
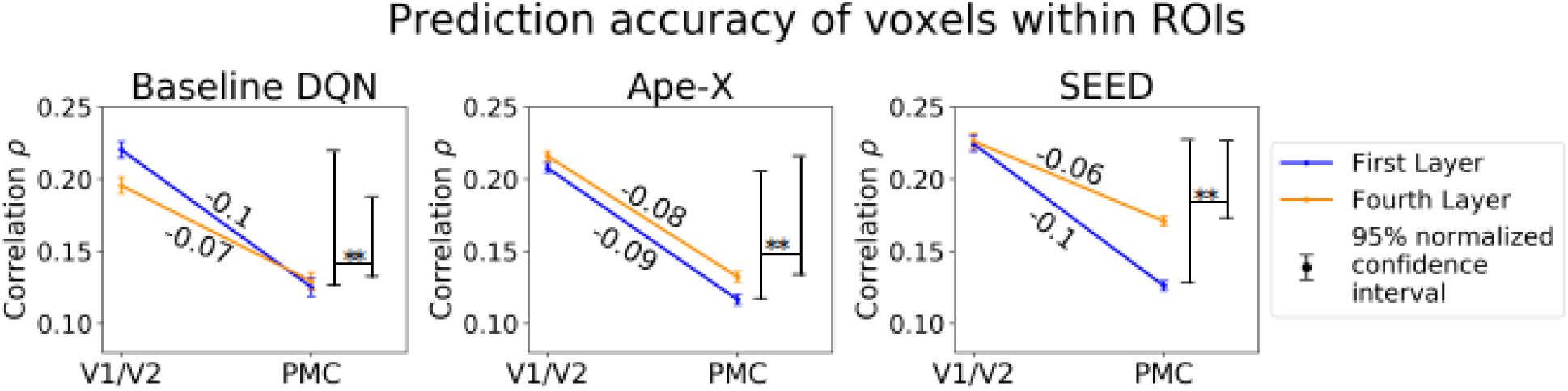
Hierarchical correspondence between the layers of a DQN and the stages of visuomotor processing. Pearson correlations were computed for voxels within V1/V2 and the PMC using features from the first and fourth layers of each DQN as predictors in the encoding model. Error bars represent the normalized 95% confidence interval of the mean correlation across subjects. The plotted lines are labeled with the gradient *ρ*(PMC) - *ρ*(V1/V2). A significant interaction between layer number and ROI was observed, reflected in an increase in the gradient with higher layer number. This increase is indicated by ‘**’ (one-tailed paired t-test, p *<* 0.01). While prediction accuracy decreases from V1/V2 to PMC for all three DQNs, the decline was less pronounced in the fourth layer than in the first, with SEED showing the most prominent increse in the gradient (repeated-measures ANOVA, *F_DQN_* (1.75) = 76, 01, Greenhouse-Geisser corrected and post hoc test, p*<* 0.01, Bonferroni-corrected). These findings, consistent with Section 3.3, provide preliminary evidence supporting a hierarchical correspondence between DQN layers and visuomotor processing stages in the human brain.

**Figure S4.**
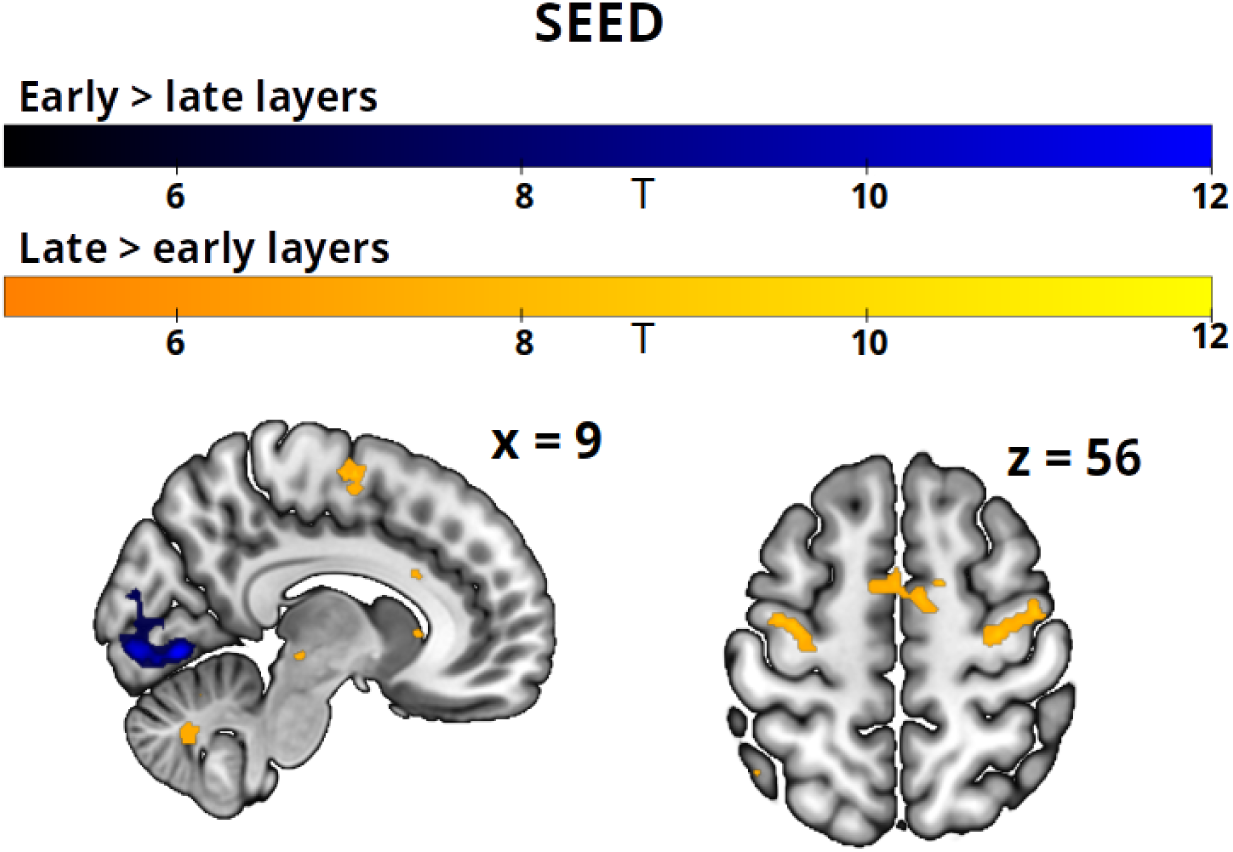
Contrasting prediction performance of early vs late layers of SEED. Voxel-wise prediction accuracies were compared between early (first, second, and third) layers and late (LSTM, fourth, and output) layers of SEED. The contrasts [1, 1, 1, −1, −1, −1] (early *>* late, shown in blue) and [−1, −1, −1, 1, 1, 1] (late *>* early, shown in orange) highlight voxels with significantly higher prediction accuracy in early and late layers, respectively. Color intensity represents the T-values of significant voxels (FWE-corrected, *p <* 0.05). Later layers provided more accurate predictions of voxel activity in higher-level PMC regions, whereas earlier layers more effectively captured voxel responses in V1/ V2.

**Figure S5.**
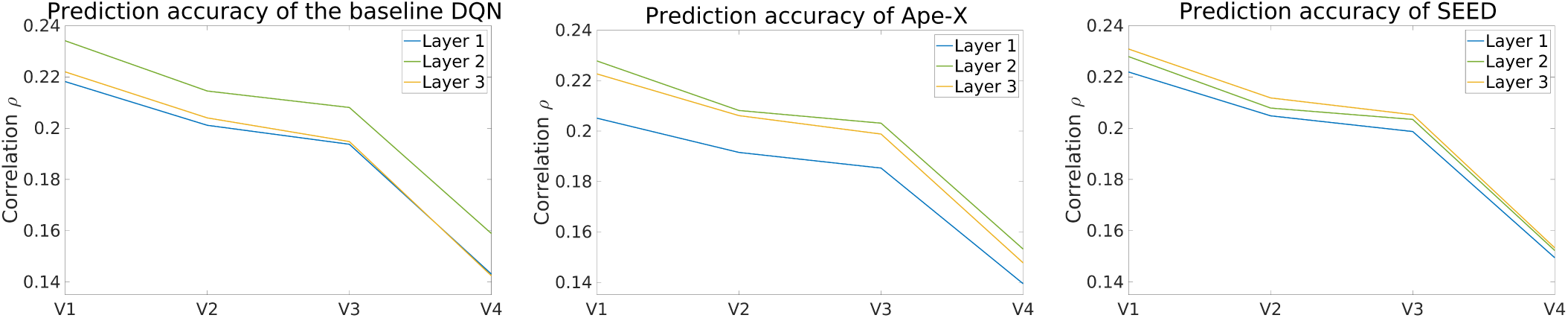
No hierarchical mapping between DQN layers and visual areas. The lines show the Pearson correlation for voxels in the visual areas V1, V2, V3, and V4. Features from the first (blue), second (green), and third (orange) convolutional layers of the DQNs were used as predictors in the encoding model. No clear hierarchical correspondence was found between the convolutional layers and the individual visual areas.

**Figure S6.**
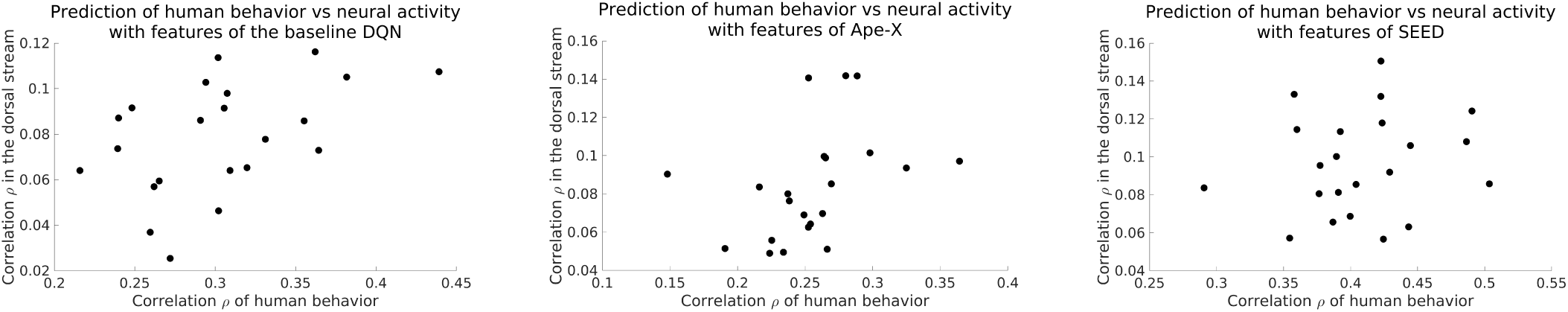
Relationship between behavioral and neural prediction accuracy. Each point represents a subject (N = 22), showing the average Pearson correlation for predicting human behavior in the form of motor responses and voxel-level neural activity along the dorsal stream. For each DQN, neurons from all layers were used as predictors. A positive relationship between behavior and neural activity was observed in the encoding models using features from the baseline DQN (*ρ* = 0.49), Ape-X (*ρ* = 0.28), and to a lesser extent, SEED (*ρ* = 0.11). These results suggest that while a model’s output may align well with behavioral data (see Supplementary Figure S11), this does not necessarily imply that the internal transformations from input to output closely mirror the underlying biological mechanisms.

**Figure S7.**
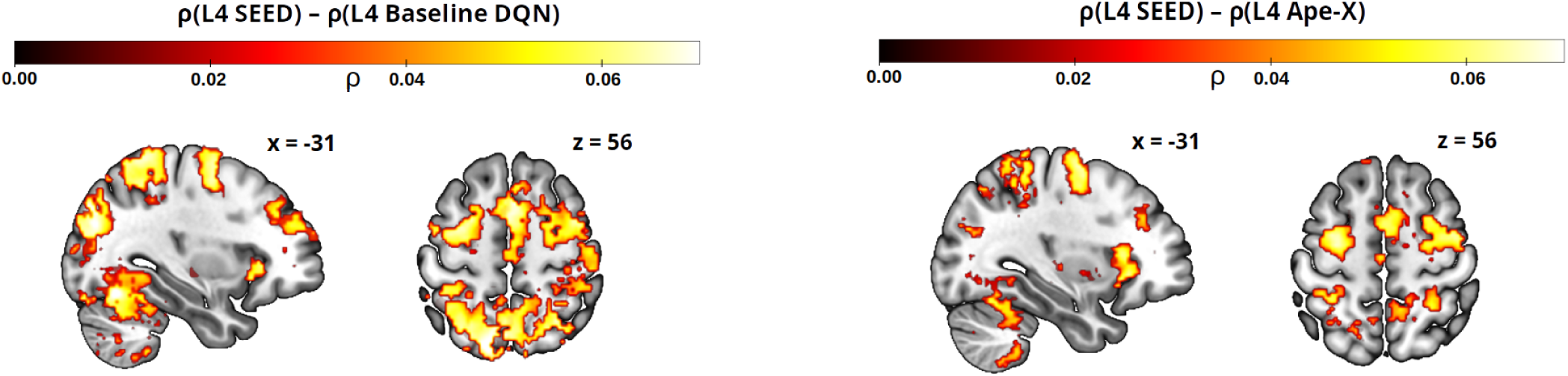
Differences in prediction accuracy by the fourth layer of the DQNs. Following the approach in Figure 5, this figure shows significant differences in Pearson correlation between actual and predicted neural activity (FWE-corrected, p *<* 0.05), computed using features from the fourth layer of SEED and those from the fourth layer of the baseline DQN (left), and those from SEED’s fourth layer and the fourth layer of Ape-X (right). As shown in Section 3.2, significant differences in prediction accuracy among the DQN features became more pronounced in higher-level control regions. Features from SEED’s fourth layer, the post-LSTM layer, more accurately predict activity in higher-level visual regions, including parts of the PPC and the PMC, compared to the fourth layers of the baseline DQN and Ape-X. These findings suggest that the LSTM may play a critical role. Its temporally integrated representations appear to enhance the model’s ability to capture brain-related features, particularly in regions associated with higher cognitive control.

**Figure S8.**
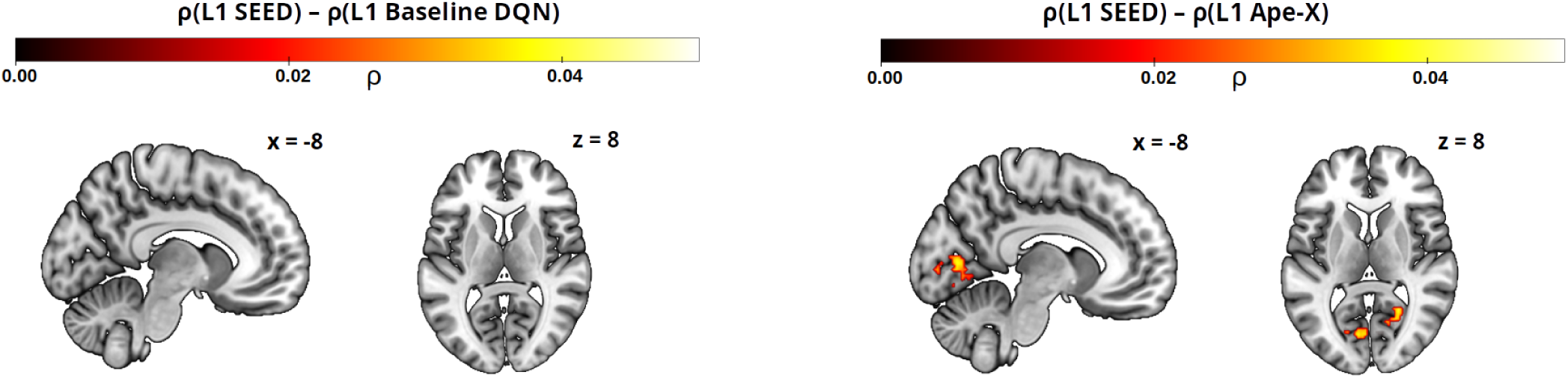
Differences in prediction of voxel activity by the first layer of the DQNs. Analogous to Supplementary Figure S7. Differences in Pearson correlation when predicting voxel activity using features from the first layer of SEED and those from the first layer of the baseline DQN (left), and from SEED’s first layer and the first layer of Ape-X (right) (FWE-corrected, p *<* 0.05). This comparison highlights that the first convolutional layers generate similar low-level features. These features span a comparable feature space across the DQNs, regardless of the subsequent architecture, leading to similar prediction performance.

**Figure S9.**
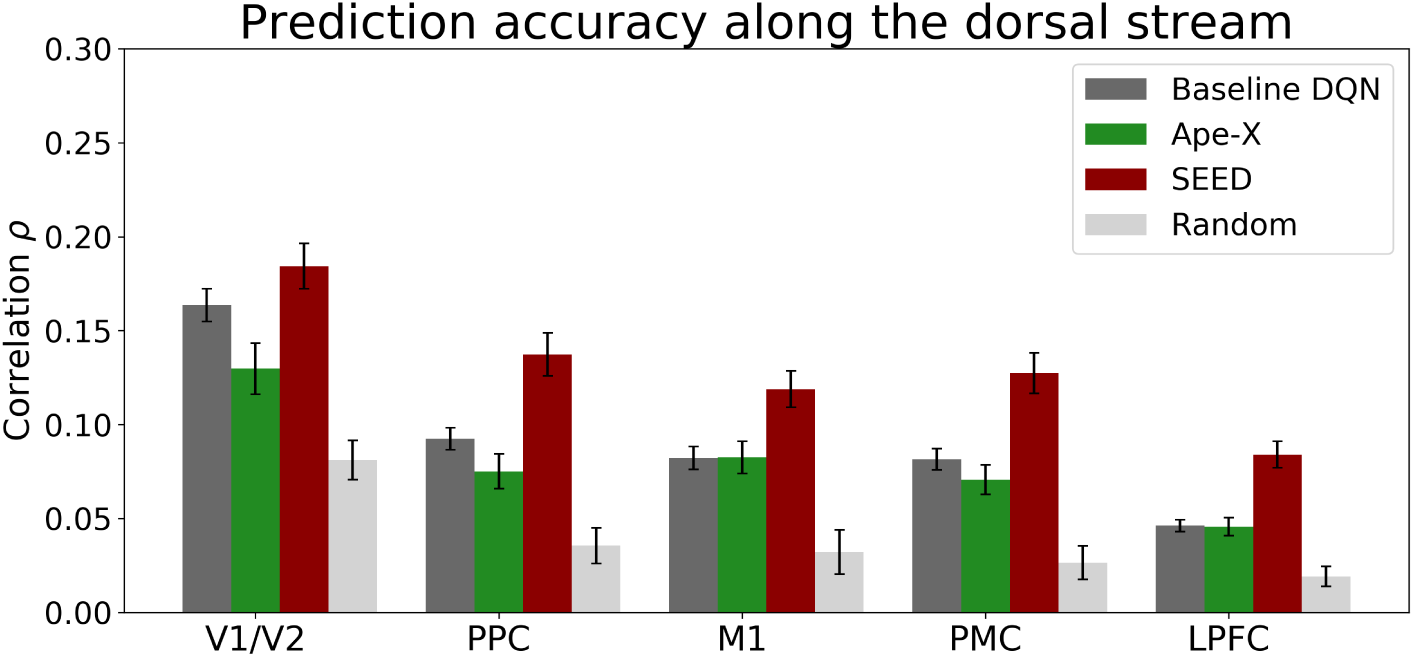
The role of training in developing effective feature generating mappings for neural prediction. For an untrained network, we used the architecture of the baseline DQN with randomly initialized weights, using three independent random initializations (light gray) on Breakout. We compared its predictive performance to that of the trained baseline DQN (dark gray), Ape-X (green), and SEED (red) in Breakout. Neurons from all layers were used as predictors in the GLM. Prediction performance was evaluated across V1/V2, PPC, M1, PMC, and LPFC. Error bars indicate the normalized 95% confidence interval of the mean Pearson correlation across subjects. As expected, the untrained DQN performed significantly worse (p *<* 0.001, Bonferroni-corrected), confirming that task-specific training is essential for extracting brain-relevant features and capturing functional representations during gameplay. Interestingly, even untrained networks with randomly initialized weights achieved above-chance prediction accuracy. This suggests that the architecture of a DQN itself, independent of its learned representations, contains inductive biases that align to some extent with features relevant for predicting neural responses. Similar findings have been reported in the same gameplay context, where DQN-generated features were used to model human behavior (Haberland et al., 2025), and in the domain of object recognition, where untrained DNNs were shown to partially explain the temporal and spatial patterns of visual brain activity (Cichy et al., 2016).

**Figure S10.**
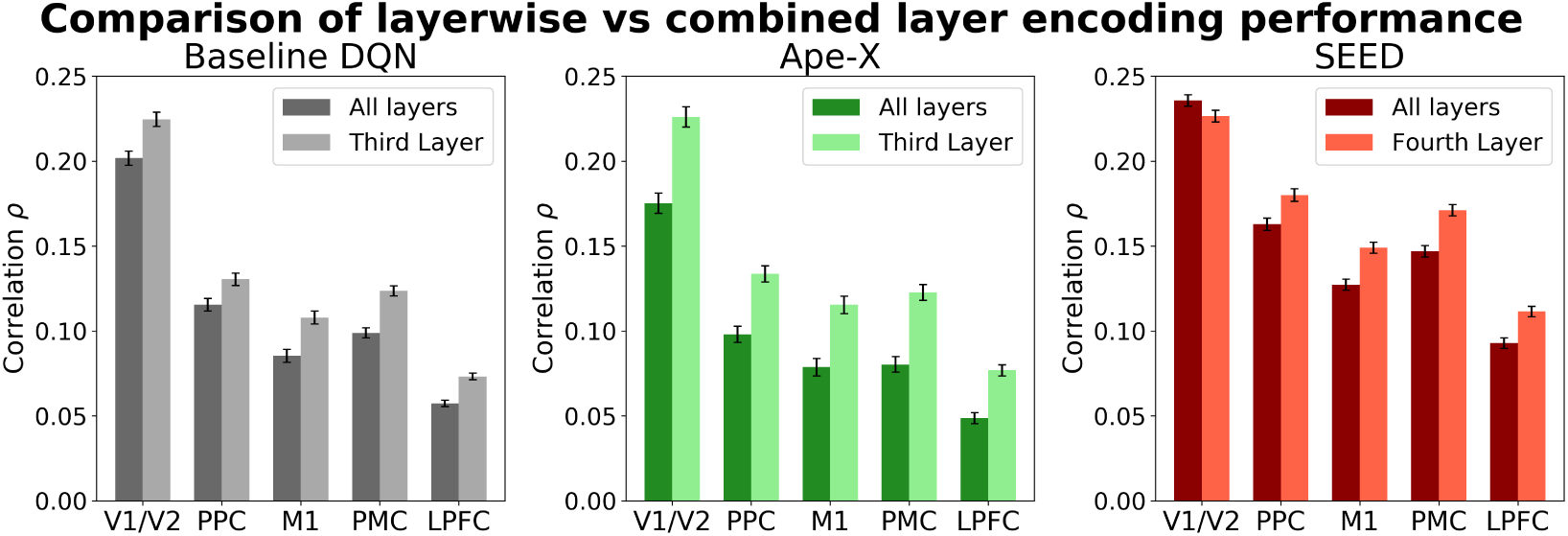
Challenges of using multiple DQN layers as predictors due to multicollinearity effects. We compared the prediction accuracy between layer-wise GLMs and GLMs using features from all layers. Bar plots show the Pearson correlation of the three encoding models in V1/V2, PPC, M1, PMC, and LPFC. Each ROI contains two bars: the left bar shows the encoding model using features from all layers of the corresponding DQN, while the right bar represents an encoding model using only features of a single specified DQN layer as predictors (the third layer of the baseline DQN and Ape-X, fourth layer of SEED). Error bars indicate the normalized 95% confidence interval of the mean Pearson correlation across subjects. Significant differences between the two GLMs were observed in all ROIs (two-sample t-test, p *<* 0.01, Bonferroni-corrected). The results highlight the impact of overfitting and multicollinearity, demonstrating that increasing the number of features in the GLM does not necessarily improve prediction performance. While this phenomenon may not hold for all layers, it is clearly observable for the ones examined in this analysis.

**Figure S11.**
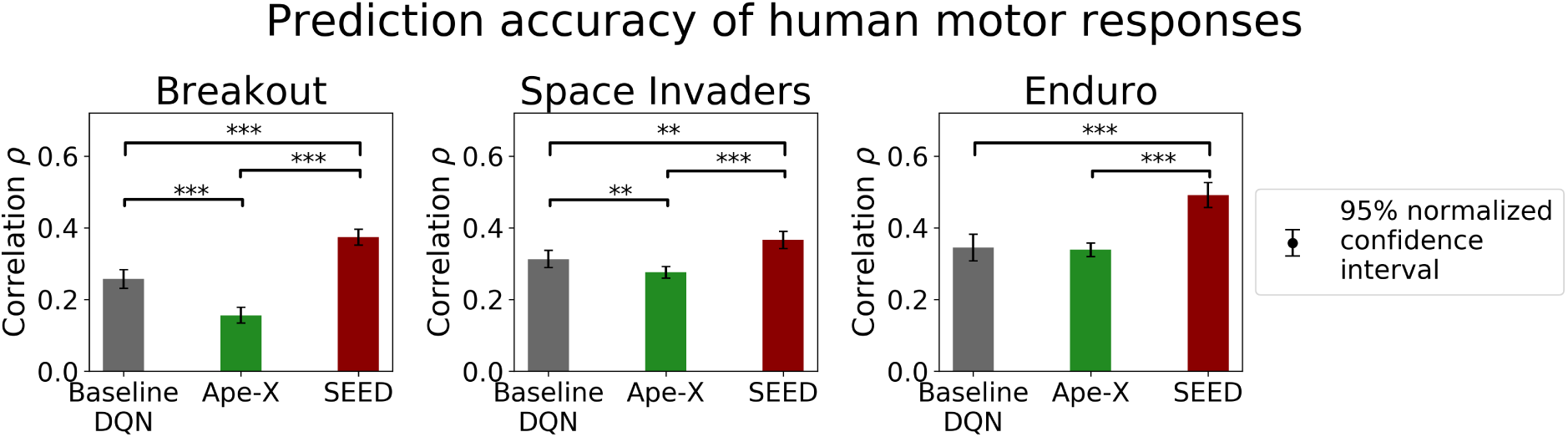
Prediction accuracy of human motor responses. Bars indicate the Pearson correlation between the actual motor responses of human subjects and the responses predicted with features of the baseline DQN (gray), Ape-X (green), and SEED (red), across Breakout (left), Space Invaders (middle), and Enduro (right). Since the output layer of a DQN contains Q-values for each possible action in a given state, these were used to predict behavior. Human behavior was predicted analogously to (Haberland et al., 2025). A smoothing kernel of FWHM of 5.3 seconds was used, consistent with the smoothing used in the fMRI data. These results align with the original behavioral study in (Haberland et al., 2025), showing that all three DQNs produce features that predict human motor responses significantly above chance level (one-sample t-test, p *<* 0.001). Among them, SEED shows the highest prediction accuracy compared to the baseline DQN and Ape-X. Significant differences are marked with ‘***’ (paired two-sample t-test, p *<* 0.001, Bonferroni-corrected) and ‘**’ (paired two-sample t-test, p *<* 0.01, Bonferroni-corrected). Error bars indicate the normalized 95% confidence interval of the mean Pearson correlation across subjects. This analysis demonstrates that the encoding models can explain not only voxel activity but also behavioral data.

**Table S1.** Effect of L1-regularization on multicollinearity among predictors. To examine the effect of L1-regularization, we assessed several metrics quantifying multicollinearity. We investigated the design matrix *A*, used to predict the fMRI time series of voxel activity, consisted of columns derived from the first layer of SEED, generated during gameplay of Breakout. To evaluate multicollinearity, we computed the number of regressors in *A*, the rank of *A*, the determinant *det*(*A^T^ A*), the condition number *κ*(*A^T^ A*), and the maximum value in the column-wise correlation matrix of *A*. We compared three different L1-regularization strengths to the case without regularization. Since GLMs were estimated voxel-wise, we randomly selected 1000 significant voxels (FWE-corrected, p *<* 0.05) and averaged the measures of multicollinearity across these voxels, as well as across the five sessions and participants. The results clearly demonstrate that increasing regularization reduces multicollinearity, as expected, highlighting the important role of regularization. In the analyses, we used a regularization parameter of *λ* = 0.07 for layer-wise prediction performance, and *λ* = 0.2 for the prediction of DQNs where features from all layers were combined as predictors in the encoding model.

| | $\lambda = 0$ | $\lambda = 0.005$ | $\lambda = 0.01$ | $\lambda = 0.07$ |
| --- | --- | --- | --- | --- |
| Number regressors | 11371.57 | 183.52 | 133.12 | 46.34 |
| Rank | 415.65 | 183.48 | 133.10 | 46.34 |
| Determinant | 0 | 4.46e+271 | 9.47e+259 | 1.05e+151 |
| Condition number | 4.99e+50 | 7.22e+15 | 1.09e+17 | 2.28e+12 |
| Correlation | 1 | 0.98 | 0.97 | 0.85 |

**Table S2.**
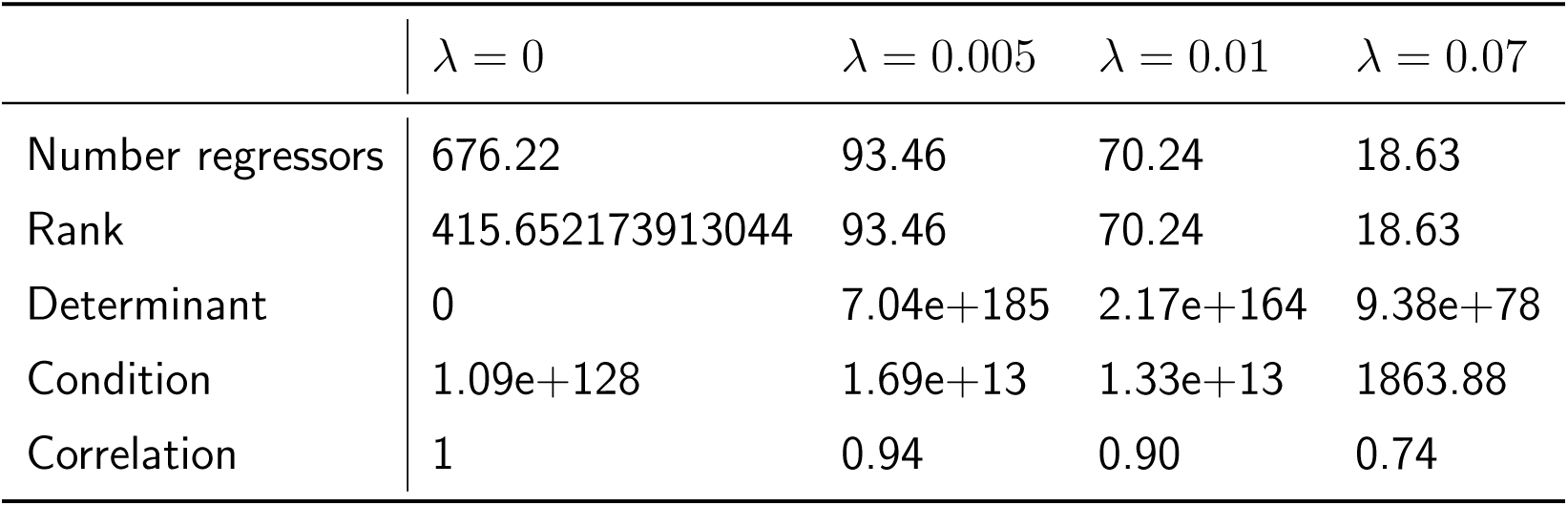
Effect of L1-regularization on multicollinearity among predictors. Same as in Supplementary Table S1 with SEED’s fourth layer.

**Table S3.** Multicollinearity between the predictors from two feature sets. To assess the dependency between predictors from two different layers, we used Canonical Correlation Analysis (CCA) to quantify the relationship between feature sets derived from the first and fourth layers of SEED during gameplay in Breakout. CCA identifies linear combinations of variables within each feature set such that the correlation between the resulting combinations is maximized, capturing the shared variance between the two sets. The table displays the top five canonical correlations for each tested regularization parameter. The analysis was conducted on 1000 randomly selected significant voxels and averaged across voxels, sessions, and participants. This analysis demonstrates that regularization effectively reduces the dependency between feature representations from different layers. It also emphasizes the inherent correlation between layers resulting from the feedforward architecture, where later layers build directly upon the representations of earlier ones. These interdependencies are particularly relevant for the GLM analyses, in which features from all layers were used simultaneously as predictors. In such cases, multicollinearity among predictors can lead to reduced model performance compared to GLMs based on features from individual layers (see Supplementary Figure S10).

| $\lambda = 0.005$ | $\lambda = 0.01$ | $\lambda = 0.07$ |
| --- | --- | --- |
| 1 | 1 | 0.97 |
| 1 | 1 | 0.95 |
| 1 | 1 | 0.93 |
| 1 | 1 | 0.92 |
| 1 | 1 | 0.84 |

**Table S4.** Custom contrast analysis for the interaction effect ROI × DQN in Figure 4.

| ROI | DQN | PPC > V1/V2 | PMC > V1/V2 | PPC > M1 | PMC > M1 |
| --- | --- | --- | --- | --- | --- |
| V1/V2 | Baseline DQN | 1 | 1 | 0 | 0 |
| PPC |  | -1 | 0 | -1 | 0 |
| M1 |  | 0 | 0 | 1 | 1 |
| PMC |  | 0 | -1 | 0 | -1 |
| LPFC |  | 0 | 0 | 0 | 0 |
| V1/V2 | Ape-X | 1 | 1 | 0 | 0 |
| PPC |  | -1 | 0 | -1 | 0 |
| M1 |  | 0 | 0 | 1 | 1 |
| PMC |  | 0 | -1 | 0 | -1 |
| LPFC |  | 0 | 0 | 0 | 0 |
| V1/V2 | SEED | -2 | -2 | 0 | 0 |
| PPC |  | 2 | 0 | 2 | 0 |
| M1 |  | 0 | 0 | -2 | -2 |
| PMC |  | 0 | 2 | 0 | 2 |
| LPFC |  | 0 | 0 | 0 | 0 |

**Table S5.** Results of the custom contrast for ROI × DQN.

| Contrast | Estimate | SE | df | t | p |
| --- | --- | --- | --- | --- | --- |
| PPC > V1/V2 | 0.015 | 0.006 | 21.000 | 2.461 | 0.023 |
| PMC > V1/V2 | 0.017 | 0.006 | 21.000 | 2.899 | 0.009 |
| PPC > M1 | 0.023 | 0.006 | 21.000 | 3.727 | 0.001 |
| PMC > M1 | 0.025 | 0.004 | 21.000 | 5.749 | < 0.001 |

**Table S6.** Custom contrast analysis for the interaction effect DQN × Layer in Figure 5.

| DQN | Layer | Differences in layer 4 > layer 1 and 3 |
| --- | --- | --- |
| Baseline DQN | Layer 1 | 0.500 |
| Ape-X |  | 0.500 |
| SEED |  | -1.000 |
| Baseline DQN | Layer 3 | 0.500 |
| Ape-X |  | 0.500 |
| SEED |  | -1.000 |
| Baseline | Layer 4 | -1.000 |
| Ape-X |  | -1.000 |
| SEED |  | 2.000 |
| Baseline DQN | Output | 0.000 |
| Ape-X |  | 0.000 |
| SEED |  | 0.000 |

**Table S7.** Results of the custom contrast for DQN × Layer.

| Contrast | Estimate | SE | df | t | p |
| --- | --- | --- | --- | --- | --- |
| 1 | 0.055 | 0.004 | 21.000 | 14.304 | < 0.001 |

**Table S8.** Custom contrast analysis for the interaction effect DQN × ROI for the fourth layer in Figure 5.

| DQN | ROI | PMC > M1 | PMC > V1/V2 | PPC > M1 | PPC > V1/V2 |
| --- | --- | --- | --- | --- | --- |
| Baseline DQN | V1/V2 | 0 | 1 | 0 | 1 |
| Ape-X |  | 0 | 1 | 0 | 1 |
| SEED |  | 0 | -2 | 0 | -2 |
| Baseline DQN | PPC | 0 | 0 | -1 | -1 |
| Ape-X |  | 0 | 0 | -1 | -1 |
| SEED |  | 0 | 0 | 2 | 2 |
| Baseline DQN | M1 | 1 | 0 | 1 | 0 |
| Ape-X |  | 1 | 0 | 1 | 0 |
| SEED |  | -2 | 0 | -2 | 0 |
| Baseline DQN | PMC | -1 | -1 | 0 | 0 |
| Ape-X |  | -1 | -1 | 0 | 0 |
| SEED |  | 2 | 2 | 0 | 0 |
| Baseline DQN | LPFC | 0 | 0 | 0 | 0 |
| Ape-X |  | 0 | 0 | 0 | 0 |
| SEED |  | 0 | 0 | 0 | 0 |

**Table S9.** Results of the custom contrast for DQN × ROI for the fourth layer.

| Contrast | Estimate | SE | df | t | p |
| --- | --- | --- | --- | --- | --- |
| PMC > M1 | 0.018 | 0.004 | 21.000 | 4.168 | < 0.001 |
| PMC > V1/V2 | 0.039 | 0.005 | 21.000 | 8.615 | < 0.001 |
| PPC > M1 | 0.025 | 0.006 | 21.000 | 4.101 | < 0.001 |
| PPC > V1/V2 | 0.046 | 0.005 | 21.000 | 8.500 | < 0.001 |

**Table S10.**
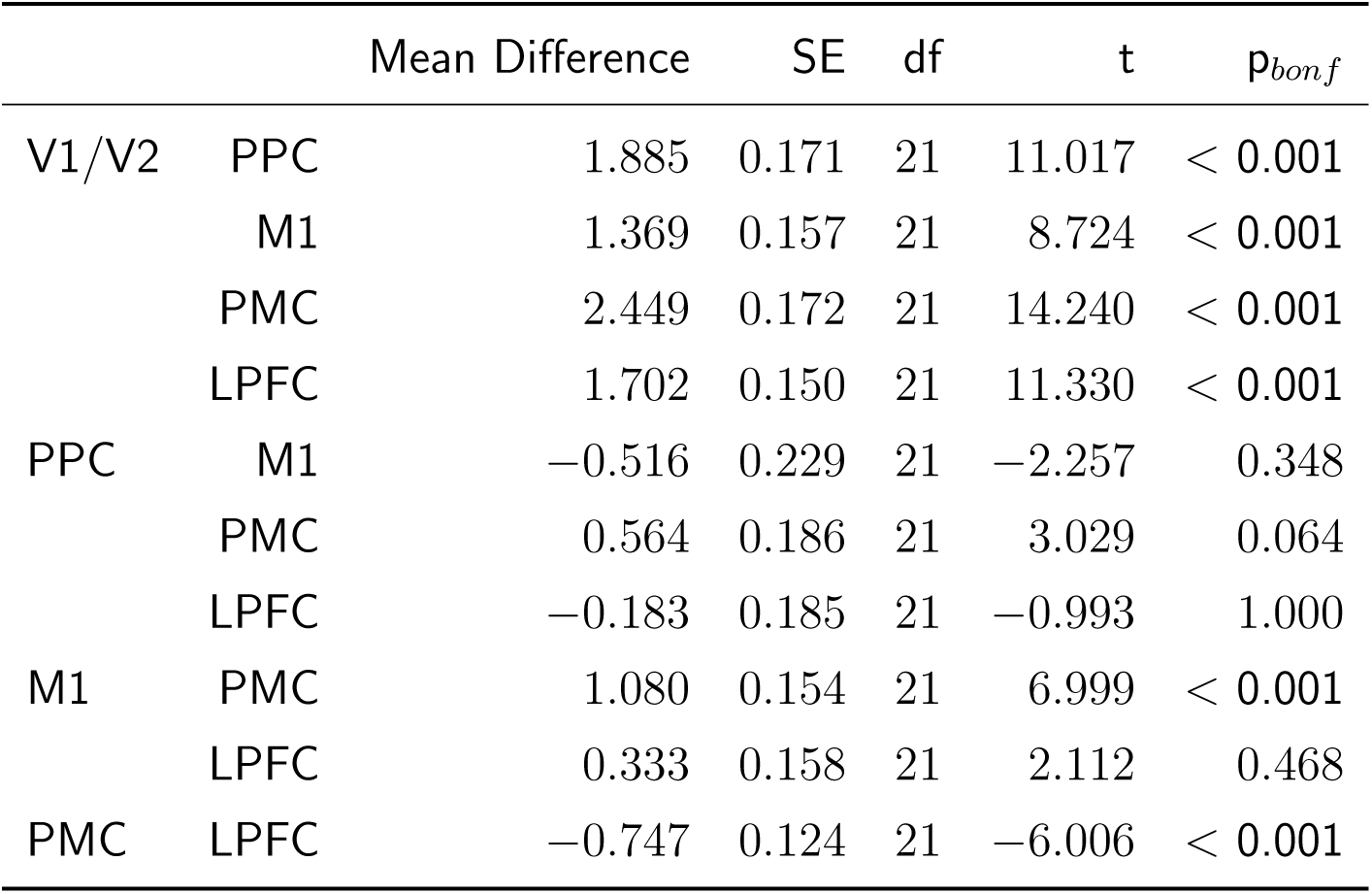
Statistical results for the comparison of early and late layer assignments in Figure 6. Post hoc comparisons for the factor *ROI*. Layer assignment analysis from Section 3.3 using the log-ratio transformation described in (Greenacre, 2021) to account for compositional data. Statistical test on the ratio of early layers (layers 1-3) to late layers (LSTM, layer 4) across the ROIs.

**Table S11.** Significant brain regions predicted by the DQNs, corresponding to Figure 2. Overview of significant cluster-level results from the SPM analysis for predicting neural activity by the encoding model using features from the baseline DQN. Reported are the corresponding brain regions (anatomical labels according to the AAL2 atlas), peak MNI coordinates (x, y, z), cluster size (k), and peak-level t-values. The p-values were FWE-corrected at the cluster-level (p *<* 0.0001) with an extent threshold of 30 voxels. Compared to the main analyses, the FWE threshold was set more conservatively (p *<* 0.0001) here for improved clarity.

| Region | x | y | z | k | t-value (peak) |
| --- | --- | --- | --- | --- | --- |
| Lingual_R | 10 | -72 | -6 | 4493 | 27.40 |
| Postcentral_R | 44 | -38 | 60 | 779 | 17.12 |
| Parietal_Sup_R | 16 | -64 | 62 | 516 | 16.90 |
| Precentral_L | -40 | -20 | 54 | 386 | 16.65 |
| Precentral_L | -32 | -6 | 60 | 169 | 13.50 |
| Insula_R | 44 | 16 | -2 | 31 | 13.47 |
| SupraMarginal_R | 58 | -42 | 42 | 47 | 13.13 |
| Temporal_Mid_L | -50 | -70 | 0 | 103 | 13.07 |
| SupraMarginal_R | 58 | -30 | 40 | 62 | 12.96 |
| Supp_Motor_Area_R | 6 | 0 | 74 | 40 | 12.86 |

**Table S12.**
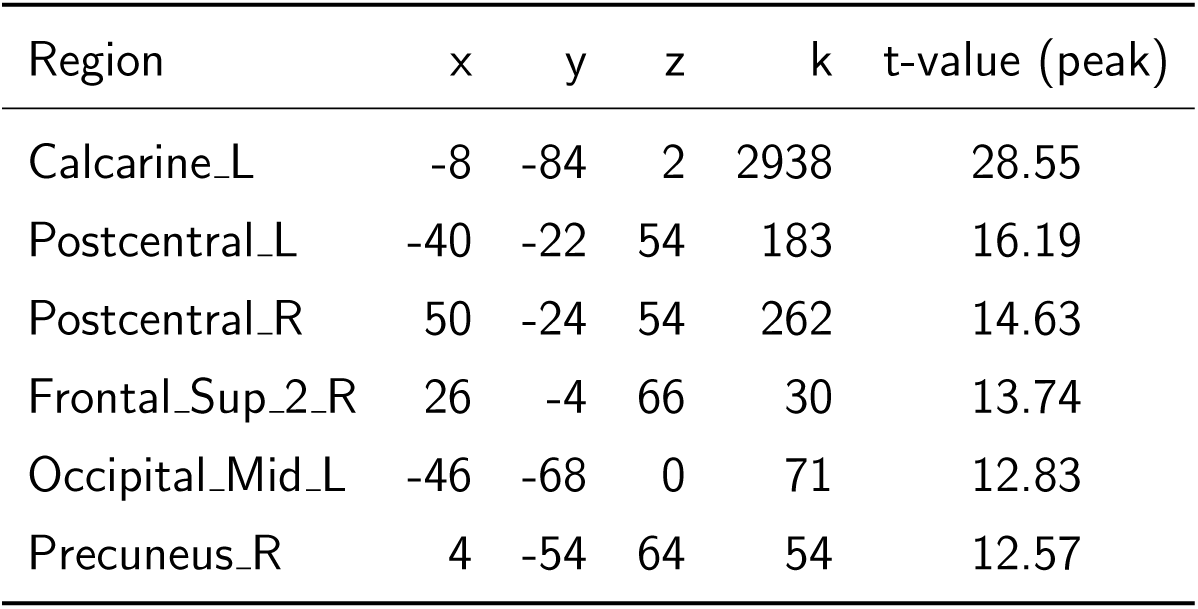
Significant brain regions predicted by the DQNs, corresponding to Figure 2. Overview of significant cluster-level results from the SPM analysis for predicting neural activity by the encoding model using features from Ape-X. Reported are the corresponding brain regions (anatomical labels according to the AAL2 atlas), peak MNI coordinates (x, y, z), cluster size (k), and peak-level t-values. The p-values were FWE-corrected at the cluster-level (p *<* 0.0001) with an extent threshold of 30 voxels. Compared to the main analyses, the FWE threshold was set more conservatively (p *<* 0.0001) here for improved clarity.

**Table S13.** Significant brain regions predicted by the DQNs, corresponding to Figure 2. Overview of significant cluster-level results from the SPM analysis for predicting neural activity by the encoding model using features from SEED. Reported are the corresponding brain regions (anatomical labels according to the AAL2 atlas), peak MNI coordinates (x, y, z), cluster size (k), and peak-level t-values. The p-values were FWE-corrected at the cluster-level (p *<* 0.0001) with an extent threshold of 50 voxels. Compared to the main analyses, the FWE threshold was set more conservatively (p *<* 0.0001) here for improved clarity.

| Region | x | y | z | k | t-value (peak) |
| --- | --- | --- | --- | --- | --- |
| Lingual_R | 10 | -72 | -8 | 24366 | 31.67 |
| Insula_R | 34 | 20 | 6 | 485 | 17.69 |
| Angular_R | 56 | -60 | 34 | 65 | 16.40 |
| Precentral_R | 58 | 8 | 36 | 241 | 16.29 |
| Angular_R | -52 | -66 | 36 | 136 | 16.24 |
| Cingulate_Mid_L | -10 | -26 | 40 | 78 | 15.93 |
| Frontal_Sup_Medial_L | -10 | 34 | 58 | 71 | 15.89 |
| Thalamus_R | 10 | -12 | 10 | 143 | 14.37 |
| Frontal_Inf_Orb_2_L | -48 | 40 | -8 | 59 | 14.26 |
| Cerebellum_9_L | -18 | -42 | -48 | 51 | 14.12 |
| Frontal_Sup_2_R | 16 | 50 | 38 | 180 | 14.07 |
| Putamen_L | -24 | 8 | 4 | 142 | 13.32 |
| Cerebellum_9_L | -8 | -56 | -56 | 96 | 13.26 |
| Frontal_Inf_Tri_R | 46 | 38 | 2 | 53 | 13.03 |

**Table S14.** Significant brain regions with differences between encoding models, corresponding to Figure 4. Overview of significant cluster-level results from the SPM analysis for differences in prediction accuracy between the encoding models using features from the baseline DQN and Ape-X. Reported are the corresponding brain regions (anatomical labels according to the AAL2 atlas), peak MNI coordinates (x, y, z), cluster size (k), and peak-level t-values. The p-values were FWE-corrected at the cluster-level (p *<* 0.05) with an extent threshold of 5 voxels.

| Region | x | y | z | k | t-value (peak) |
| --- | --- | --- | --- | --- | --- |
| Frontal_Sup_2_L | -24 | -8 | 64 | 7 | 8.41 |
| Occipital_Mid_L | -24 | -84 | 26 | 7 | 8.12 |

**Table S15.** Significant brain regions with differences between encoding models, corresponding to Figure 4. Overview of significant cluster-level results from the SPM analysis for differences in prediction accuracy between the encoding models using features from SEED and the baseline DQN. Reported are the corresponding brain regions (anatomical labels according to the AAL2 atlas), peak MNI coordinates (x, y, z), cluster size (k), and peak-level t-values. The p-values were FWE-corrected at the cluster-level (p *<* 0.05) with an extent threshold of 50 voxels.

| Region | x | y | z | k | t-value (peak) |
| --- | --- | --- | --- | --- | --- |
| Supp_Motor_Area_R | 4 | 8 | 56 | 22230 | 21.68 |
| Frontal_Inf_Tri_L | -36 | 20 | 8 | 439 | 13.96 |
| Insula_R | 34 | 26 | -4 | 1422 | 13.61 |
| Caudate_L | -16 | 0 | 20 | 652 | 13.35 |
| Frontal_Sup_2_R | 24 | 56 | 28 | 192 | 13.09 |
| Postcentral_L | -62 | -12 | 30 | 394 | 12.26 |
| Cerebellum_8_R | 36 | -48 | 52 | 55 | 11.64 |
| Temporal_Mid_R | 62 | -38 | -8 | 50 | 10.62 |
| Cerebellum_8_R | 16 | -66 | -46 | 83 | 10.26 |
| Frontal_Sup_2_L | -26 | 34 | 28 | 63 | 10.06 |
| Parietal_Inf_L | -44 | -50 | 38 | 73 | 9.84 |

**Table S16.**
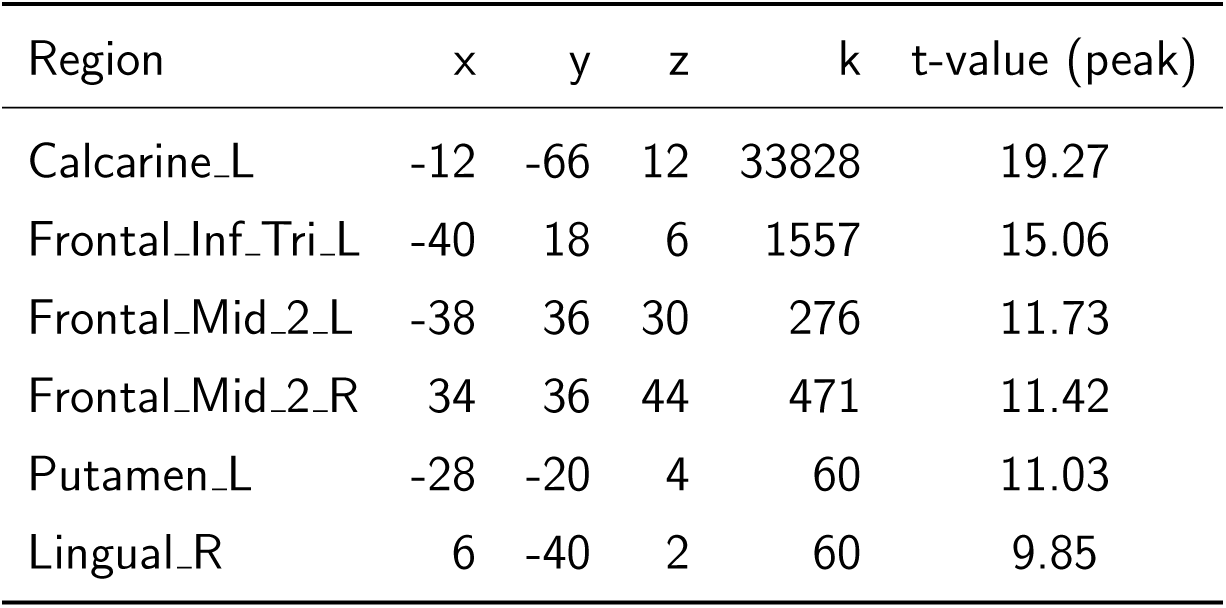
Significant brain regions with differences between encoding models, corresponding to Figure 4. Overview of significant cluster-level results from the SPM analysis for differences in prediction accuracy between the encoding models using features from SEED and Ape-X. Reported are the corresponding brain regions (anatomical labels according to the AAL2 atlas), peak MNI coordinates (x, y, z), cluster size (k), and peak-level t-values. The p-values were FWE-corrected at the cluster-level (p *<* 0.05) with an extent threshold of 50 voxels.

| Region | x | y | z | k | t-value (peak) |
| --- | --- | --- | --- | --- | --- |
| Calcarine_L | -12 | -66 | 12 | 33828 | 19.27 |
| Frontal_Inf_Tri_L | -40 | 18 | 6 | 1557 | 15.06 |
| Frontal_Mid_2_L | -38 | 36 | 30 | 276 | 11.73 |
| Frontal_Mid_2_R | 34 | 36 | 44 | 471 | 11.42 |
| Putamen_L | -28 | -20 | 4 | 60 | 11.03 |
| Lingual_R | 6 | -40 | 2 | 60 | 9.85 |

